# Sex differences in the alcohol-mediated modulation of BLA network states

**DOI:** 10.1101/2022.01.07.475435

**Authors:** Alyssa DiLeo, Pantelis Antonodiou, Spencer Ha, Jamie L. Maguire

## Abstract

About 85% of adults in the United States report drinking alcohol in their lifetime. Mood disorders, like generalized anxiety disorder and major depression, are highly comorbid with alcohol use. The basolateral amygdala (BLA) is an area of the brain that is heavily implicated in both mood disorders and alcohol use disorder. Importantly, modulation of BLA network/oscillatory states via parvalbumin-positive (PV) GABAergic interneurons has been shown to control the behavioral expression of fear and anxiety. Further, PV interneurons express a high density of δ-subunit-containing GABA_A_ receptors (GABA_A_Rs), which are sensitive to low concentrations of alcohol. Our lab previously demonstrated that δ-subunit-containing GABA_A_Rs on PV interneurons in the BLA influence voluntary ethanol intake and anxiety-like behavior in withdrawal. Therefore, we hypothesized that the effects of alcohol may modulate BLA network states that have been associated with fear and anxiety behaviors via δ-GABA_A_Rs on PV interneurons in the BLA. Given the impact of ovarian hormones on the expression of δ-GABA_A_Rs, we examined the ability of alcohol to modulate local field potentials (LFPs) in the BLA from male and female C57BL/6J and *Gabrd^-/-^* mice after acute and repeated exposure to alcohol. Here, we demonstrate that acute and repeated alcohol can differentially modulate oscillatory states in male and female C57BL/6J mice, a process which involves δ-GABA_A_Rs. This is the first study to demonstrate that alcohol is capable of altering network states implicated in both anxiety and alcohol use disorders.

**Significance Statement:** Alcohol use disorder and mood disorders are highly comorbid. The basolateral amygdala (BLA) is implicated in both of these disorders, but the mechanisms contributing to their shared pathophysiology remain uncertain. Here we demonstrate that acute and repeated alcohol exposure can alter network oscillations in the BLA which control the behavioral expression of fear and anxiety. These data suggest that alcohol may directly influence network states associated with mood. Further, we demonstrate sex differences in alcohol’s ability to modulate BLA network states, an effect involving δ-GABA_A_ receptors, which may contribute to sex differences in alcohol intake and comorbid mood disorders. These data potentially point to a novel mechanism mediating the effects of alcohol on affective states.

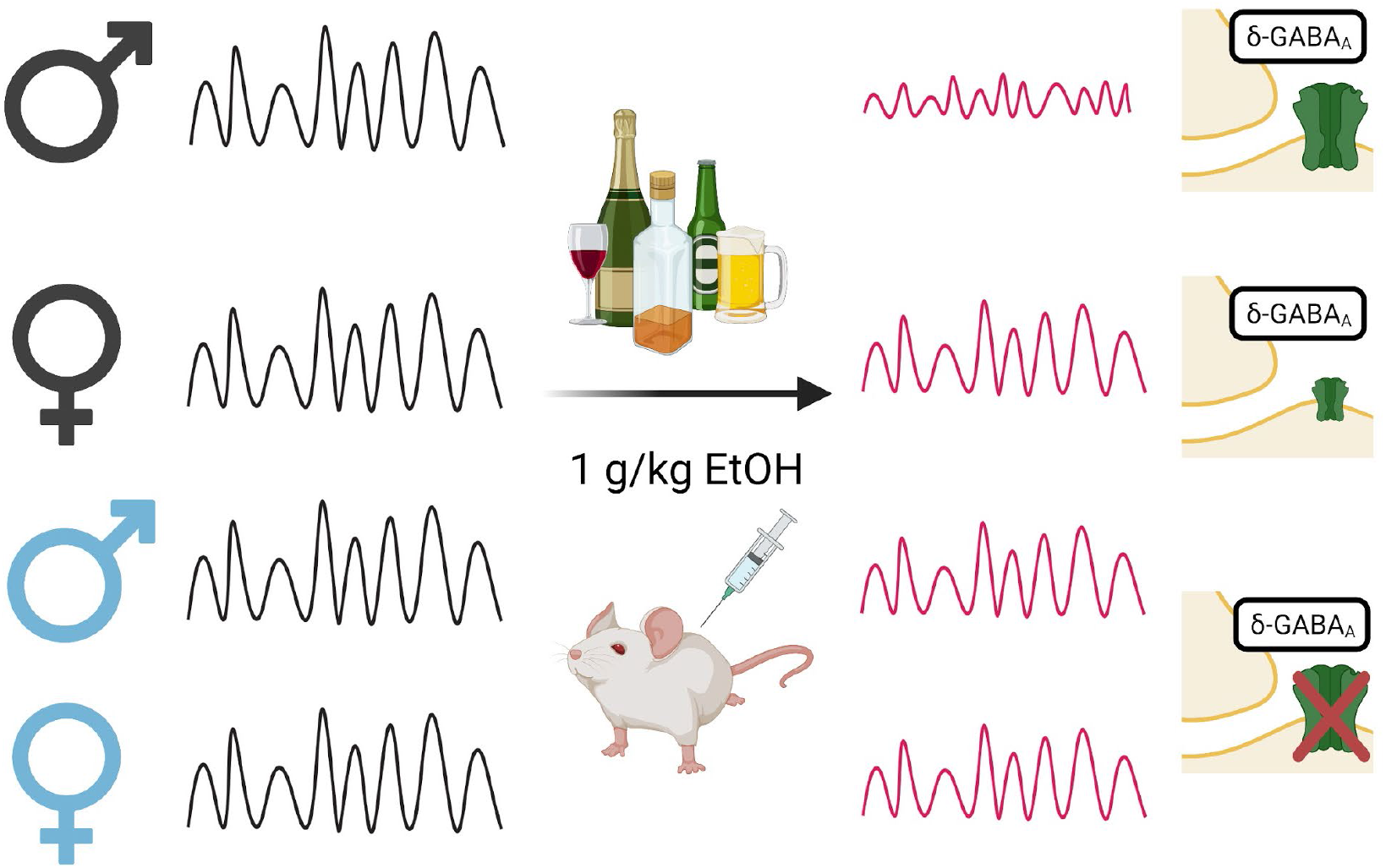

## Introduction

Alcohol is the most widely used drug in the United States, with approximately 85% of adults reporting alcohol use in their lifetime. Despite this high rate of use, only about 5% will go on to develop an alcohol use disorder while most adults continue to drink without reaching this diagnostic criterion (SAMHSA, 2020). The transition from first drink to alcohol dependence is encouraged by both the positive and negative reinforcing effects of alcohol, each with corresponding neurobiological frameworks (Gilpin and Koob, 2008). Comorbid mood disorders, such as major depression and anxiety disorders, contribute to the reinforcing effects of alcohol by pushing individuals to drink to relieve tension in high stress or high anxiety situations (Kushner et al., 2011). The basolateral amygdala (BLA) has been identified as a brain region at the intersection of mood disorders and alcohol, contributing to both alcohol use disorder and anxiety disorders (Silberman et al., 2009; Tye et al., 2011; Agoglia and Herman, 2018).

Accumulating evidence demonstrates a critical role for oscillatory states in the BLA in modulating fear and anxiety states (Likhtik et al., 2013; Stujenske et al., 2014; Davis et al., 2017; Antonoudiou et al., 2021). However, the impact of alcohol on these network states has not been explored. Network oscillations within and between brain areas represent a mechanism for transition between brain and behavioral states. Specifically, reciprocal connections between the BLA and the medial prefrontal cortex (mPFC) have been shown to be important for fear expression and anxiety-like behaviors (Likhtik et al., 2013). Our lab has recently demonstrated that particular oscillation frequencies within and between the BLA and mPFC are associated with either a fear (3-6 Hz) or safety (6-12 Hz) state (Davis et al., 2017). Further, this circuit, along with other regions like the hippocampus, has also been shown to contribute to high and low anxiety states in mice (Likhtik et al., 2013).

It is well established that the anxiolytic properties of alcohol can motivate consumption and contribute to the high comorbidity of alcohol use disorders and mood disorders (Thomas et al., 2003; Smith and Randall, 2012; Mason et al., 2018). However, it is unclear how alcohol impacts network states underlying modulation of anxiety states. Here we examine the ability of acute, low dose alcohol to modulate BLA network activity in alcohol naïve mice, using local field potentials (LFPs) to measure network oscillations in the BLA in male and female C57BL/6J mice during acute and repeated exposure to alcohol.

The generation of oscillations is thought to involve the ability of GABAergic interneurons, particularly parvalbumin (PV) expressing interneurons, to synchronize populations of principal neurons (Bartos et al., 2007; Sohal et al., 2009; Fuchs et al., 2017). Somatic-targeting, fastspiking PV interneurons exert powerful control over a large network of excitatory principal cells, and as such, they are capable of synchronizing the network and generating oscillations to orchestrate network communication (Mcdonald, 1992; Bocchio and Capogna, 2014). There is a critical role for PV interneurons in oscillation generation within the BLA both *ex vivo* and *in vivo,* where PV interneurons can shift oscillatory frequencies and drive behavioral states (Antonoudiou et al., 2021; Ozawa et al., 2020; Davis et al., 2017).

Our lab has also demonstrated that PV interneurons in the BLA express a high density of extrasynaptic δ subunit-containing GABA_A_Rs, which are uniquely sensitive to alcohol and play a role in regulating both alcohol consumption and anxiety-like behaviors, including anxiety associated with alcohol withdrawal (Glykys et al., 2007; Melón et al., 2018; Antonoudiou et al., 2021). Tonic inhibition mediated by δ-GABA_A_Rs has been shown to control hippocampal oscillations (Mann and Mody, 2010; Pavlov et al., 2014) and loss of the δ subunit in PV interneurons alters gamma oscillations in the CA3 region of the hippocampus (Ferando and Mody, 2013, 2014). Given the evidence that PV interneurons modulate oscillations in the BLA and their high density of δ subunit expression, we further hypothesized that alcohol acts through δ-GABA_A_Rs in the BLA to modulate oscillations associated with the network communication of fear and anxiety. To test this, we examined the ability of alcohol to alter oscillatory states in the BLA of male and female *Gabrd^-/-^* mice. Our findings suggest that the ability of alcohol to modulate network states involves δ subunit-containing GABA_A_Rs. We conclude that alcohol can modulate BLA oscillatory states in a sex-specific manner, a process which, in part, involves δ subunit-containing GABA_A_Rs.

## Materials and Methods

### Animals

Adult male and female C57BL/6J mice, aged 8-12 weeks old, were purchased from Jackson Lab (stock #000664) and group housed at Tufts University School of Medicine in temperature and humidity-controlled housing rooms on a 12-hour light-dark cycle (lights on at 7AM) with ad libitum food and water. Animals were handled according to protocols and procedures approved by the Tufts University Institutional Animal Care and Use Committee (IACUC). Global *Gabrd^-/-^* knockout mice were bred in house (Mihalek et al., 1999, 2001). Mice were single housed and habituated to new cages for 24 hours before the start of experiments.

### Stereotaxic Surgery

All mice undergoing surgery were anesthetized with ketamine/xylazine (90 mg/kg and 5-10 mg/kg, respectively, i.p.) and treated with sustained release buprenorphine (0.5-1.0 mg/kg, s.c.). A lengthwise incision was made to expose the skull and a unilateral craniotomy was performed to lower a depth electrode (PFA-coated stainless-steel wire, A-M systems) into the BLA (AP - 1.50 mm, ML 3.30 mm, DV – 5 mm), affixed to a head mount (Pinnacle #8201) with stainless steel screws as ground, reference, and frontal cortex EEG (AP +0.75 mm, ML ± 0.3 mm, DV −2.1 mm) electrodes. EMG wires were placed in the back of the neck.

### LFP Recordings

LFP recordings were performed as previously described (Antonoudiou et al., 2021) in male and female C57BL/6J and *Gabrd-^-/-^* mice after a week of recovery from implant surgery. LFP recordings were acquired using Lab Chart software (AD Instruments) collected at 4 KHz and amplified 100X. The data were band pass filtered (1-300 Hz) and spectral analysis was performed in MATLAB (Antonoudiou et al., 2021) using MatWAND (https://github.com/pantelisantonoudiou/MatWAND) which utilizes the fast Fourier transform similar to previous reports (Kruse and Eckhorn, 1996; Frigo and Johnson, 1998; Pape et al., 1998; Freeman et al., 2000). Briefly, recordings were divided into 5 second overlapping segments and the power spectral density for a range of frequencies was obtained (Oppenheim et al., 1999).

### Acute & Repeated Alcohol Exposure

Mice were habituated to new cages with ad libitum food and water for 24 hours before starting the experimental paradigm. All injections were performed 2-3 hours into the light cycle at the same time each day across all cohorts. The acute exposure consisted of a 60-minute baseline period followed by a saline injection (0.9% NaCl i.p.) and a subsequent 1 g/kg ethanol injection (20% v/v i.p.). The repeated exposure consisted of a 60 minute baseline period followed by an i.p. injection of either saline or 1 g/kg ethanol (20% v/v) daily for five consecutive days. For the females, the acute ethanol exposure was calculated from the first day of the repeated exposure paradigm.

### Immunohistochemistry

Immunohistochemistry was performed as previously reported (Melón et al., 2018) in a separate cohort of mice following the acute or repeated alcohol exposure paradigms. Mice were anesthetized with isoflurane, transcradially perfused with 0.9% saline and 4% paraformaldehyde (PFA), the brains were rapidly excised, fixed in 4% PFA overnight, and subsequently cryoprotected in 10% and 30% sucrose. The brains were then flash frozen using isopentane and stored at −80°C until cryosectioning. Free floating 40 μm coronal slices were co-stained for PV and δ using universal antigen retrieval buffer (R&D systems) and primary antibodies against δ-GABA_A_R (1:100, Phosphosolutions) and PV (1:1000, Sigma P3088) for 72 hours at 4°C. The slices were then incubated with a biotinylated goat anti-rabbit (1:1000, Vector Laboratories BA1000) and Alexa-Fluor 647 conjugated goat anti-mouse (1:200, ThermoFisher Scientific A28181) for two hours at room temperature and streptavidin conjugated Alexa-Fluor 488 (1:200, ThermoFisher Scientific S32354) for two hours at room temperature. Slices were mounted and cover slipped with antifade hard set mounting medium with DAPI (Vectashield H1500). Fluorescent labeling in the basolateral amygdala was imaged on a Nikon A1R confocal microscope and z-stacks were acquired using a 20X objective. Camera settings were kept consistent across samples and cohorts. The images were analyzed using Image J software by outlining PV-positive interneurons using the ROI manager and measuring the integrated density of PV and δ expression on the outlined PV-positive interneurons. Each cell was considered its own data point within each animal.

### Statistical Analysis

Data were analyzed using Prism 8 software (GraphPad) and MatWAND in MATLAB (Mathworks) as previously described (Antonoudiou et al., 2021). To ensure a consistent time period for analysis across cohorts, we analyzed the first 40 minutes of baseline and the first 35 minutes of each injection period. Repeated measures two-way ANOVAs were performed to detect significance of frequency, treatment, sex, or genotype. A Greenhouse-Geisser correction was applied where necessary. A mixed effects model was used if values were missing across days. A post-hoc Šídák’s multiple comparisons test was performed to identify significant differences of specific frequency ranges. ANOVA results are reported in Extended Data Table 1-1 and 4-1. Multiple comparisons are reported in Extended Data Table 1-2 and 4-2. P values < than 0.05 were considered significant. All *n* values for each treatment group are shown in the figure legends.

## Results

### Alcohol modulates BLA network states

To characterize the effect of acute alcohol on BLA oscillations in wild type mice, we recorded LFPs in the BLA of male C57BL/6J mice in response to either vehicle (0.9% saline i.p) or alcohol (1 g/kg i.p) injection (Figure 1A, B). We found that the vehicle injections significantly decreased the power area of BLA oscillations high theta (6-12 Hz) (p = 0.0117, 95% C.I = [0.04658, 0.4029]), low gamma (40-70 Hz) (p = 0.0027, 95% C.I = [−0.9799, −0.2066]), and high gamma (80-120 Hz) range (p = 0.0103, 95% C.I = [−0.6491, −0.08079]) as compared to baseline (Figure 1C). However, we did not find any difference between the two vehicle injections, indicating there was no sensitization or adaptation to the second injection. The impact of vehicle injections on oscillatory states in the BLA has been observed previously (Antonoudiou et al., 2021) and is thought to reflect the network response to the stress of the injection. Therefore, all results are compared to the first vehicle injection (Figure 1B).

**Figure 1.**
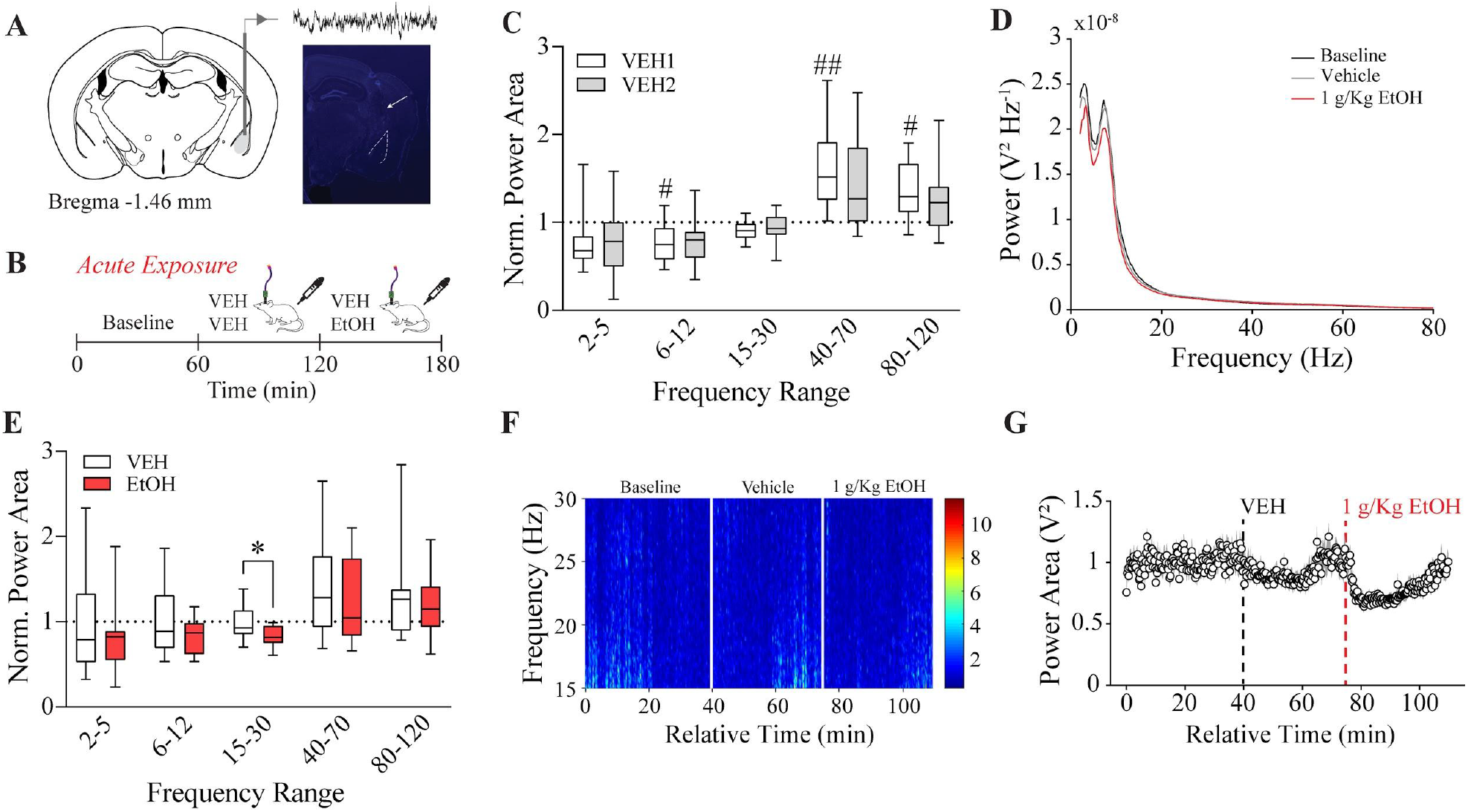
Acute alcohol alters BLA network activity in male C57BL/6J mice. **A,** Representative targeting of local field potential recordings from the basolateral amygdala. **B,** Acute alcohol exposure paradigm consisted of local field potential recordings during baseline (60 minutes), vehicle (0.9% saline i.p.; 60 minutes) injection, and a treatment injection (0.9% saline or 1 g/kg alcohol i.p.; 60 minutes). **C,** Normalized power area for vehicle/vehicle acute exposure (*n* = 15). **D,** Power spectral density of baseline, vehicle, and 1 g/kg alcohol injection over 0-80 Hz. **E,** Normalized power area for vehicle/alcohol acute exposure (*n* = 14). **F,** Representative spectrogram of normalized BLA LFP beta power (15-30 Hz) from vehicle/alcohol acute exposure experiment. **G,** Average normalized beta power area (15-30 Hz) during acute injections of vehicle and alcohol (1 g/kg i.p). Dots represent the mean and the shaded region represents SEM. #p < 0.05, ##p < 0.01 vs. baseline, *p < 0.05 vs. vehicle.

In response to alcohol treatment in male mice, the power area in the beta frequency (15-30 Hz) is decreased compared to vehicle (p = 0.034, 95% C.I = [0.01033, 0.2925]; Figure 1D-G). This indicates that alcohol can modulate specific oscillatory frequencies within the BLA that are implicated both in addiction and mood disorders (Jurado-Barba et al., 2020).

### Alcohol modulates BLA network states in a sex-dependent manner

Because of the well documented sex differences in alcohol related behaviors (Melón et al., 2013; Barkley-Levenson and Crabbe, 2015; Becker and Koob, 2016; Sneddon et al., 2019), we treated female C57BL/6J to the same acute alcohol paradigm as described for the males (Figure 1B). Similar to the males, we did not find any significant differences between the two vehicle injections in the vehicle/vehicle control experiments in females (Figure 2-1A). We did find that vehicle significantly decreased high theta BLA LFP power area as compared to baseline (p = 0.0273, 95% C.I = [0.2699, 0.4296]; Figure 2A), similar to what we observed in the males. Additionally, there was no significant difference between the male and female C57BL/6J BLA LFP response to the vehicle injection (Figure 2-1B).

**Figure 2.**
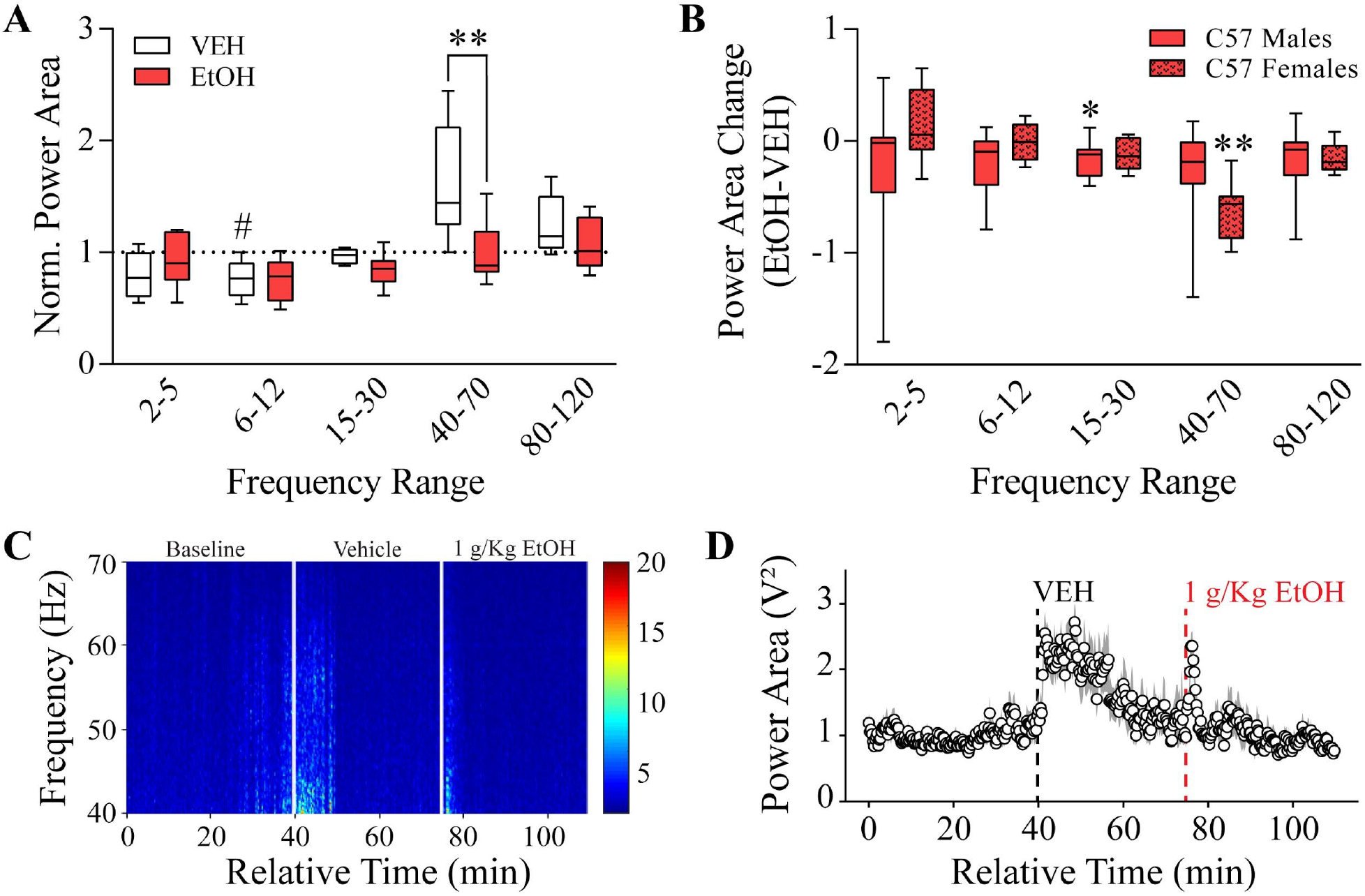
Acute alcohol alters different BLA oscillatory frequencies in female C57BL/6J mice. **A,** Normalized power area for vehicle/alcohol acute exposure (*n* = 8). **B,** Power area difference between the vehicle and alcohol injection period of C57BL/6J male and female mice. **C,** Representative spectrogram of normalized BLA LFP gamma power (40-70 Hz) from vehicle/alcohol acute exposure experiment in female C57BL/6J mice. **D,** Average normalized gamma power area (40-70 Hz) during acute injections of vehicle and alcohol (1 g/kg i.p) in female C57BL/6J mice. Dots represent the mean and the shaded region represents SEM. #p < 0.05 vs. baseline, *p < 0.05, **p < 0.01 vs. vehicle.

In response to acute alcohol exposure (EtOH-vehicle), we found that alcohol significantly decreased the gamma band BLA LFP power area as compared to vehicle in female C57BL/6J mice (p = 0.0014, 95% C.I = [0.2955, 0.9441]; Figure 2A, C, D), a unique signature from the males. Although alcohol decreased BLA LFP power area in different frequency bands in males and females, there were no direct significant differences between groups (Figure 2B).

Collectively, these data suggest that acute ethanol modulates the BLA network differently in male and female mice.

### Alcohol modulation of BLA network states involves δ subunit-containing GABA_A_Rs

Previous literature has supported the role of δ-GABA_A_Rs in mediating the effects of alcohol on tonic inhibition, drinking and withdrawal behaviors (Wallner et al., 2003; Santhakumar et al., 2007; Melón et al., 2018; Darnieder et al., 2019). Therefore, to test the role of this receptor subunit on mediating the ability of alcohol to modulate BLA network states, we repeated the same procedure as described above in male and female *Gabrd*^-/-^ mice. We found that vehicle injections significantly increased BLA LFP power area at low gamma frequencies only in the vehicle/alcohol condition in male *Gabrd^-/-^* mice as compared to baseline (p = 0.0121, 95% C.I = [−1.062, −0.1299]; Figure 3A). In both *Gabrd^-/-^* males and females, we did not find any significant difference between vehicle injections (Figure 3-1A, B).

**Figure 3.**
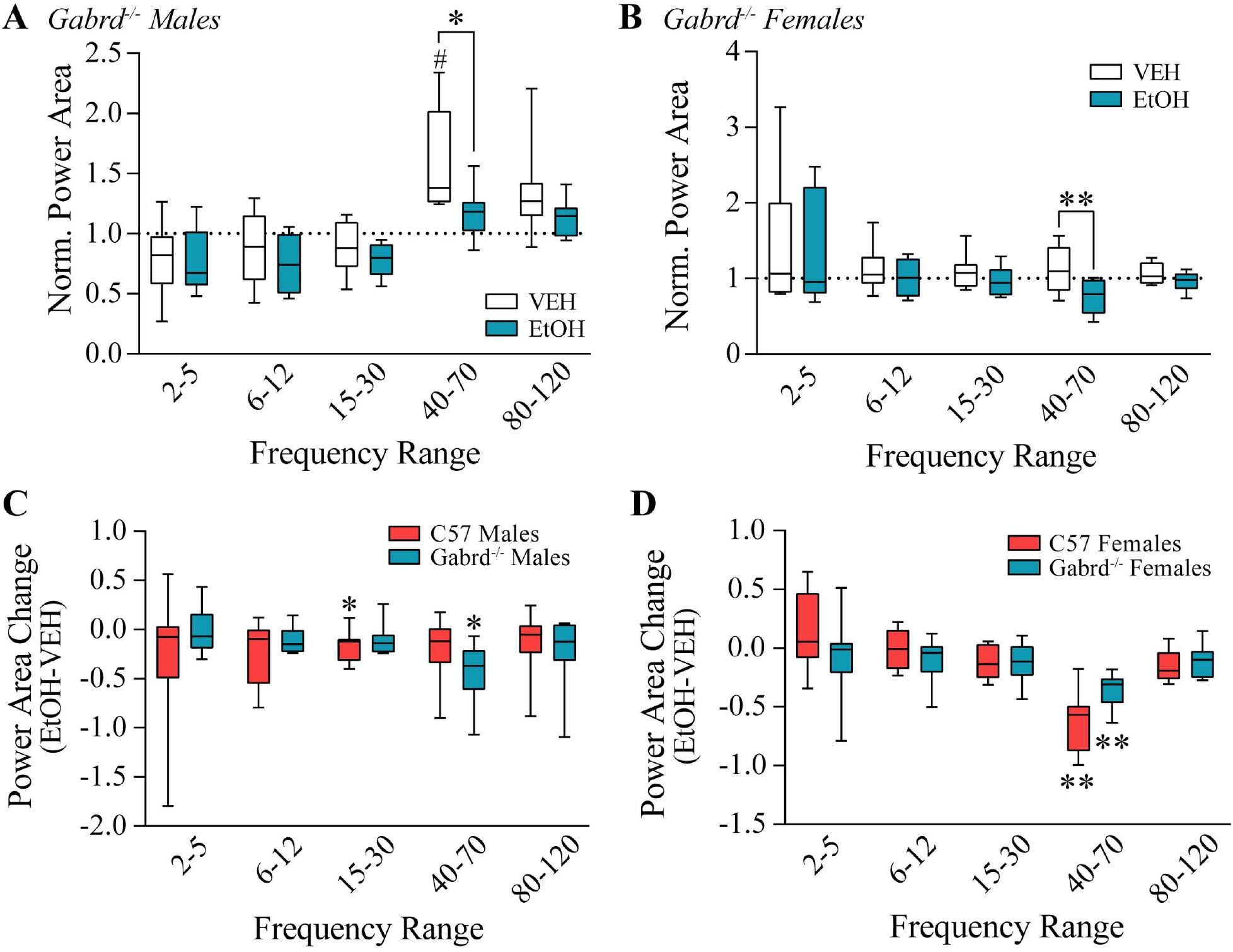
Acute alcohol in Gabrd^-/-^ mice decreases the same BLA frequency bands as female C57BL/6J mice. **A,** Normalized power area for vehicle/alcohol acute exposure in male Gabrd^-/-^ mice (*n* = 9) and **B,** female Gabrd^-/-^ mice (*n* = 8). **C,** Power area difference between the vehicle and alcohol injection period of male C57BL/6J and Gabrd^-/-^ mice. **D,** Power area difference between the vehicle and alcohol injection period of female C57BL/6J and Gabrd^-/-^ mice. #p < 0.05, ##p < 0.01 vs. baseline, *p < 0.05, **p < 0.01 vs. vehicle.

Unlike C57BL/6J males, acute alcohol significantly decreased the low gamma band of males (p = 0.020, 95% C.I = [0.07, 0.78]; Figure 3A) and females (p = 0.0012, 95% C.I = [0.1731, 0.5352]; Figure 3B) as compared to vehicle. This effect was similar to, but not as robust an effect, as in C57BL/6J females. However, direct comparisons did not detect significant differences in the ability of alcohol to modulate oscillatory states between male C57BL/6J and male *Gabrd^-/-^* mice (Figure 3C) or between C57BL/6J females and *Gabrd^-/-^* females (Figure 3D). Collectively, these data suggest that the impact of the loss of the GABA_A_R δ subunit on the effect of alcohol are more profound in males and induces similar effects as observed in C57BL/6J females.

### Ability of repeated alcohol exposure to modulate BLA network states is dependent on δ subunit-containing GABA_A_Rs

Since we established that acute alcohol could modulate specific oscillatory frequencies in the BLA, we were interested in how BLA LFPs changed over time in response to repeated doses of alcohol. Male C57BL/6J and *Gabrd^-/-^* mice received vehicle (0.9% saline) or low dose (1 g/kg i.p.) alcohol for five consecutive days (Figure 4A). We did not find significant effects of repeated vehicle injections within baseline (Figure 4B top) or as an effect of vehicle (Figure 4B bottom) across days in either the male C57BL/6J or male *Gabrd^-/-^* mice (Figure 4-1).

**Figure 4.**
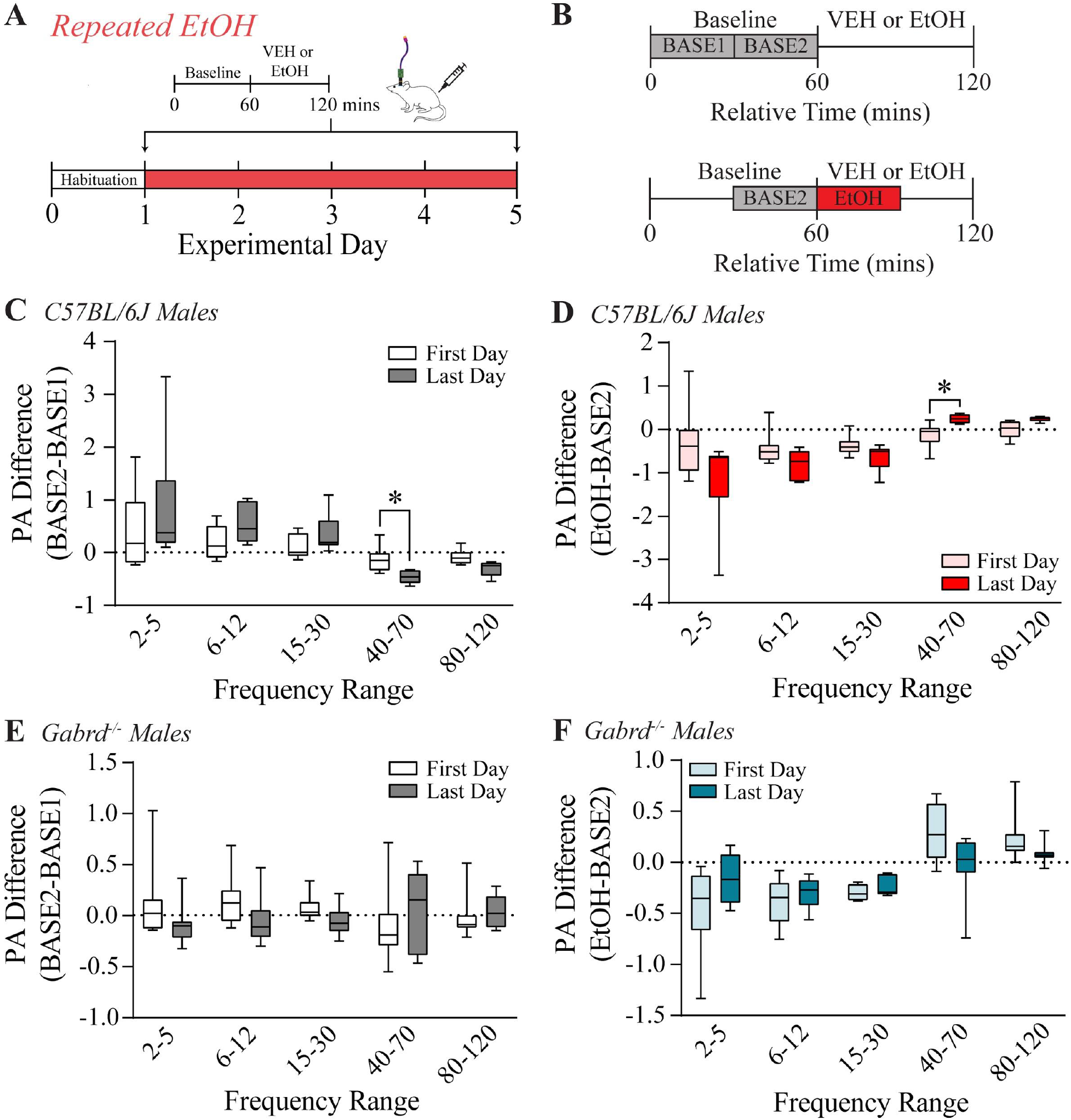
Repeated exposures of alcohol further decreases BLA LFP oscillations in male C57BL/6J mice. **A,** Experimental paradigm of the repeated alcohol exposure procedure in male C57BL/6J and Gabrd^-/-^ mice of BLA LFP recordings during baseline (60 minutes) and vehicle or alcohol injections (0.9% saline or 1 g/kg alcohol i.p.; 60 minutes) over five days. **B,** Analysis format which measured the change in baseline (BASE2-BASE1) and change in response to vehicle or alcohol injections (VEH or EtOH - BASE2). **C,** Change in baseline on the first and last day of exposure in male C57BL/6J mice (first day *n* = 8; last day *n* = 6) and **E,** male Gabrd^-/-^ mice (first day *n* = 8; last day *n* = 8). **D,** Change in effect of alcohol on the first and last day of exposure in male C57BL/6J mice and **F,** male Gabrd^-/-^ mice. *p < 0.05 vs first exposure.

Interestingly, in response to repeated treatment, we found a change in the baseline low gamma BLA LFP power from the first to last day (BASE2-BASE1) in male C57BL/6J mice (p = 0.0252, 95% C.I = [0.03785, 0.6384]; Figure 4C), which may be an anticipatory change associated with repeated alcohol administration. In addition, in response to alcohol treatment, we observed a significant increase in low gamma BLA LFP power area from the first to last day (EtOH-BASE2) (p = 0.0305, 95% C.I = [−0.7087, −0.03272]; Figure 4D). In contrast, we did not observe significant effects of repeated alcohol on baseline or treatment in the male *Gabrd^-/-^* mice (Figure 4E, F).

Direct comparison between male C57BL/6J and *Gabrd^-/-^* mice on the first day of alcohol exposure does not reveal any significant changes within baseline (Figure 5A), but did find that male C57BL/6J mice had significantly decreased high theta (p = 0.0138, 95% C.I = [−1.061, −0.1088]) and beta (p = 0.0022, 95% C.I = [−0.7193, −0.1659]) BLA LFP power area as compared to male *Gabrd^-/-^* mice in response to alcohol exposure (Figure 5B). By the last day, there were significant decreases in the low (p = 0.0343, 95% C.I = [−0.9892, −0.03576]) and high gamma (p = 0.0062, 95% C.I = [−0.5908, −0.09553]) BLA LFP power area in male C57BL/6J mice compared to male *Gabrd^-/-^* mice (Figure 5C), again likely attributed to the role of the GABA_A_R δ subunit in the anticipatory effects of repeated alcohol exposure. In response to repeated alcohol administration, we observe a significant increase in the high gamma frequency range in male C57BL/6J mice compared to male *Gabrd^-/-^* mice (p = 0.0203, 95% C.I = [0.02211, 0.2899]; Figure 5D). Overall, these results suggest a blunted impact of acute and repeated alcohol exposure on BLA oscillatory states in mice lacking the GABA_A_R δ subunit. Further, these data indicate a role for δ-GABA_A_Rs in adapting to alcohol exposure over time, as well as anticipating alcohol treatment as shown by the changes in baseline in male C57BL/6J mice, but not *Gabrd^-/-^* mice.

**Figure 5.**
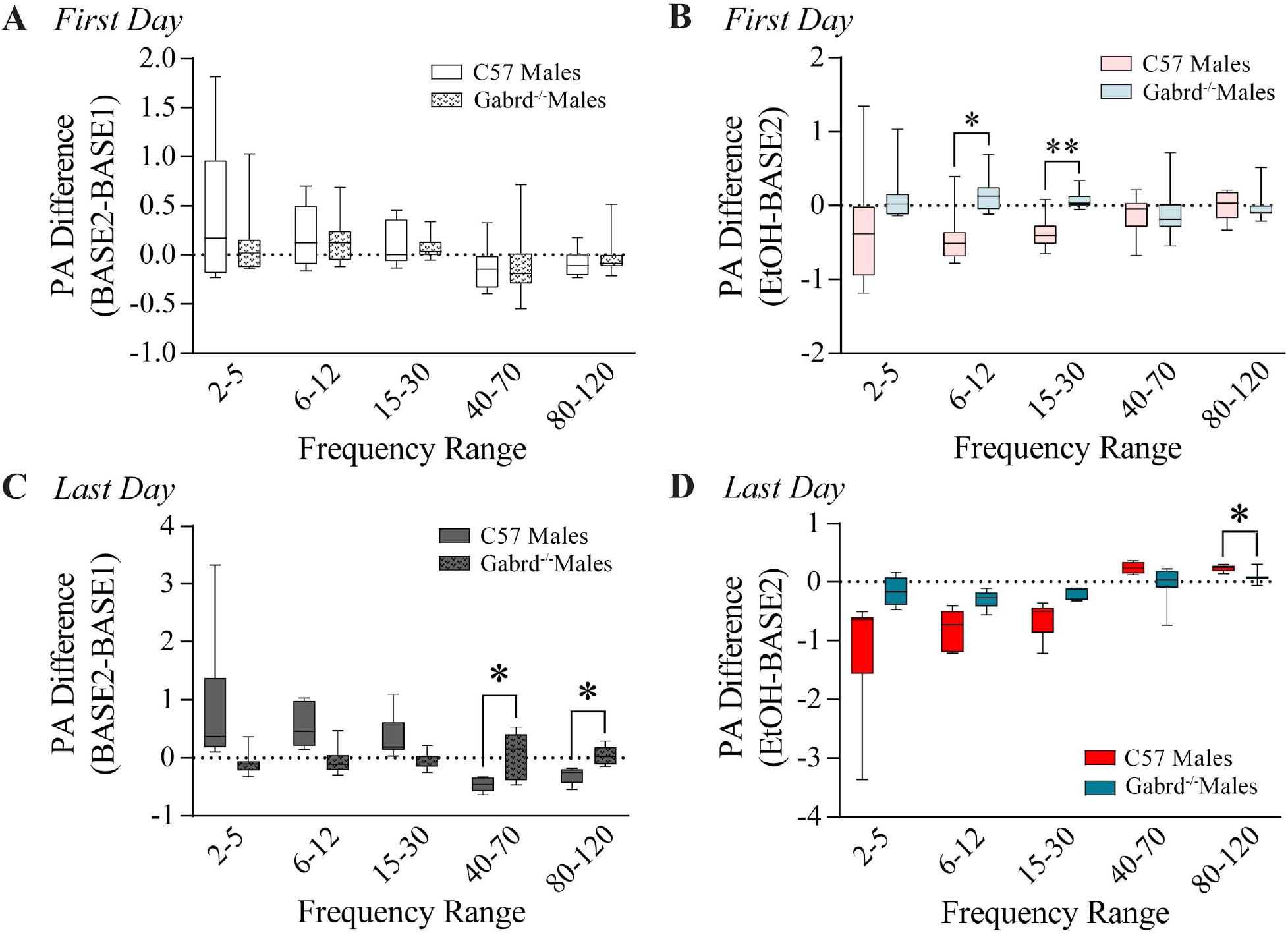
Effects of repeated alcohol BLA LFP modulation is dependent on δ subunit-containing GABA_A_Rs. **A,** Comparison between male C57BL/6J and Gabrd^-/-^ mice in the change in baseline on the first and C, last day of exposure. **B,** Comparison between male C57BL/6J and Gabrd^-/-^ mice in the change in response to alcohol on the first and D, last day of exposure. *p < 0.05, **p < 0.01 vs. Gabrd^-/-^ mice.

### Sex differences in BLA network states in response to repeated alcohol exposure

Repeated alcohol exposure in female C57BL/6J and *Gabrd^-/-^* mice involved acute alcohol or vehicle exposure on day one and the repeated alcohol exposure days two to five (Figure 6A). We will be using their second day of exposure in our repeated alcohol comparisons, which were not significantly different in female C57BL/6J mice (Figure 6B). Neither day one or day two were significantly different from day five in female *Gabrd^-/-^* mice (Figure 6-1A, B). Therefore, we continued to use day two as the first day of repeated exposure in our analysis.

**Figure 6.**
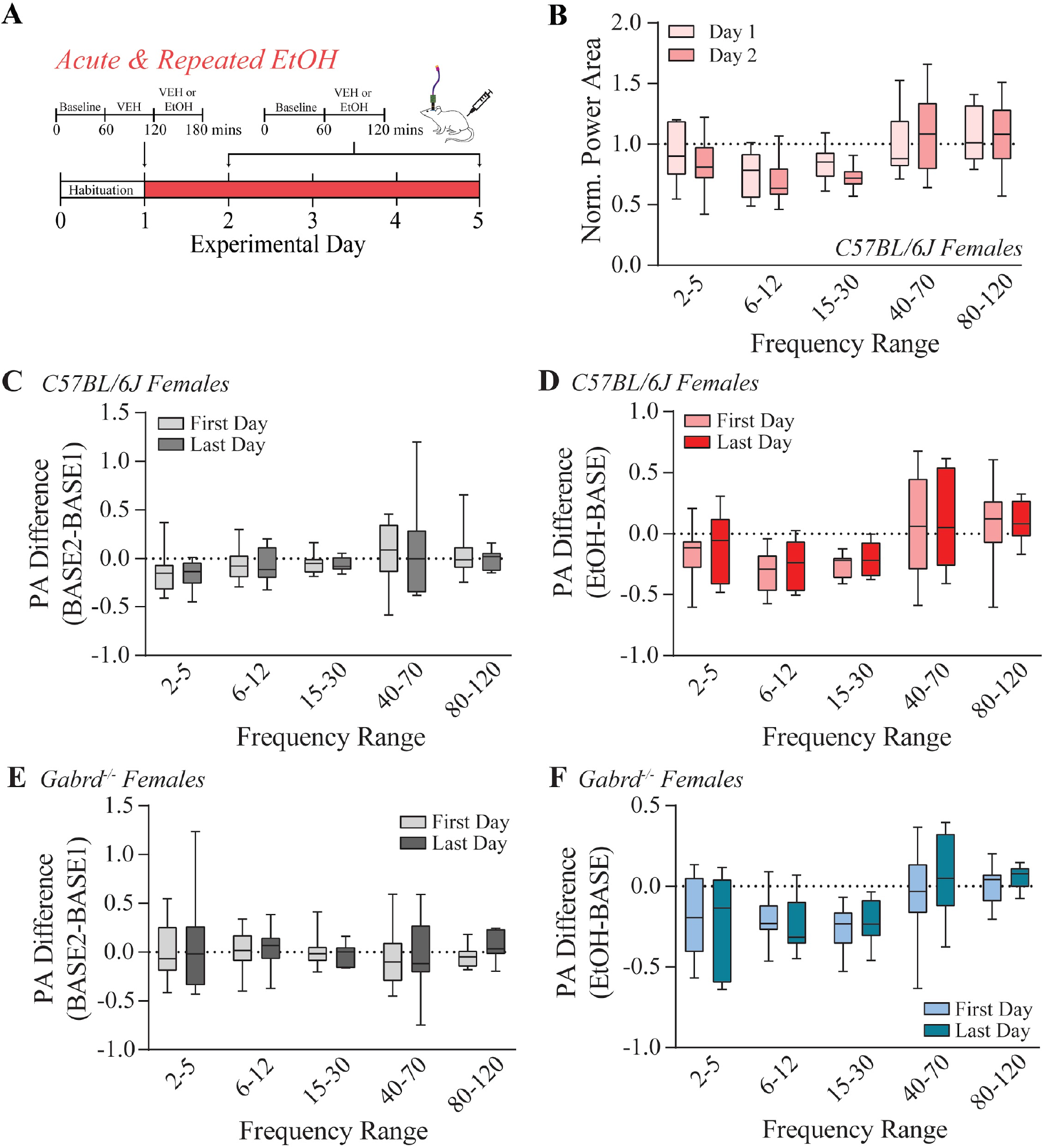
Repeated alcohol does not modulate BLA LFPs in female C57BL/6J and Gabrd^-/-^ mice. **A,** Repeated alcohol paradigm for female C57BL/6J and Gabrd^-/-^ mice which includes the acute alcohol exposure (day one) and the repeated alcohol exposure as day two (first day) to five (last day). **B,** Normalized power area of BLA LFP of alcohol injections during the acute alcohol injection (day one) and day two (first day) in female C57BL/6J mice (*n* = 10). **C,** Change in baseline on the first and last day of exposure (first day *n* = 8; last day *n* = 7). **D,** Change in effect of alcohol on the first and last day of exposure in female C57BL/6J and **F,** Gabrd^-/-^ mice.

We did not observe significant differences within the baseline or in the effect of vehicle between the first and last day of exposure in female C57BL/6J (Figure 6-2A, B) or female *Gabrd^-/-^* mice (Figure 6-2C, D). Interestingly, unlike the males, we did not observe any significant effect of repeated alcohol in either group across days (Figure 6C-F) or between the C57BL/6J female and *Gabrd^-/-^* females when directly compared (Figure 6-3A-D).

Direct comparison between C57BL/6J males and females did not reveal significant differences at any frequency range within the baseline period (Figure 7A) or in the effect of alcohol from baseline (Figure 7B) on the first day of exposure. However, by the last day, we found specific an increase in high theta (p = 0.0435, 95% C.I = [0.01858, 1.216]) and a decrease in high gamma BLA LFP power area at baseline of males with no change in females (p = 0.0122, 95% C.I = [-0.5198, −0.06395]; Figure 7C). In response repeated alcohol exposure, males exhibited a significantly reduced BLA LFP power area in the high theta frequency range in C57BL/6J males with no change in females (p = 0.0433, 95% C.I = [−1.085, −0.01596]; Figure 7D). Collectively, these data suggest that females show tolerance to repeated alcohol exposure compared to males, an effect that involves the GABA_A_R δ subunit.

**Figure 7.**
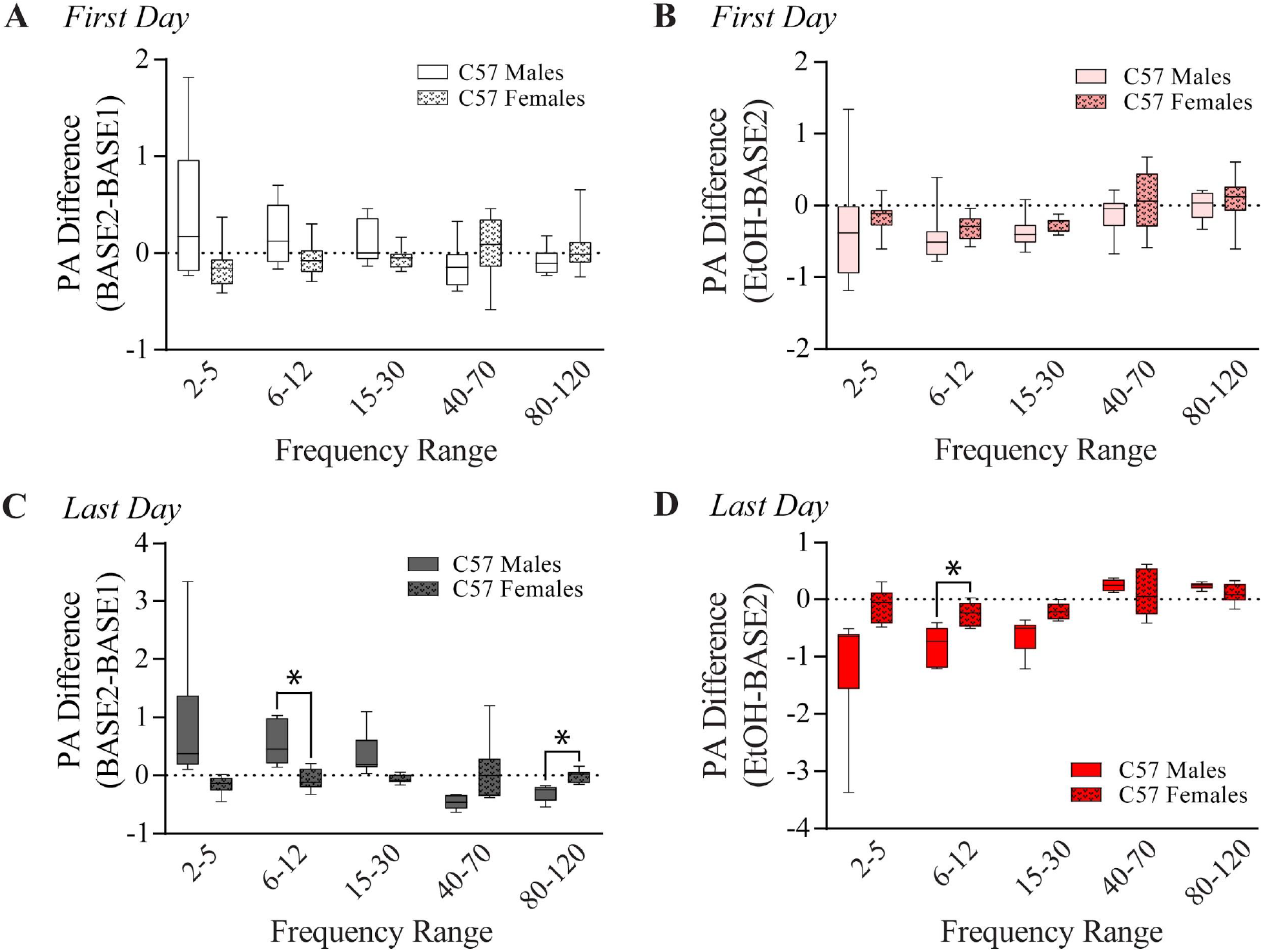
Sex differences in BLA LFP response to repeated alcohol exposure. **A,** Comparison between male and female C57BL/6J mice in the change in baseline on the first day of repeated exposure and **C,** day five. **B,** Comparison between male and female C57BL/6J mice in the change in response to alcohol on the first day of exposure and **D,** on the last day of exposure. *p < 0.05 vs. female C57BL/6J mice.

### Repeated alcohol exposure alters δ expression on PV interneurons in the BLA

Alcohol exposure can change the expression of GABA_A_R subunits (Liang et al., 2004; Olsen et al., 2012; Lindemeyer et al., 2014; Follesa et al., 2015) and sex differences in GABA_A_R δ subunit expression has been reported (Maguire et al., 2005). Given that changes in the expression of the GABA_A_R δ subunit, whether through genetic deletions or associated with hormone fluctuations during pregnancy, can alter oscillation frequencies in the hippocampus (Ferando and Mody, 2013, 2015), we hypothesized that altered expression of the GABA_A_R δ subunit, expressed on PV interneurons in the BLA, may contribute to our observed sex differences in BLA network states. Thus, we examined whether there were any potential changes sex differences in GABA_A_R δ subunit expression in the BLA or changes in expression associated with alcohol exposure. First we investigated potential sex differences in GABA_A_R δ subunit expression in the BLA. We observe an higher expression δ expression on PV interneurons in naive female C57BL/6J mice (*M* = 1002542, *SEM* = 45011) as compared to naive male C57BL/6J mice (*M* = 538252, *SEM* = 12440; t[424] = 10.39, p < 0.0001; Figure 8B) with no change to PV immunoreactivity (female: *M* = 1464808, *SEM* = 51507; male: *M* = 1448517, *SEM* = 97320; Figure 8A). Interestingly, vehicle treatment alone reduced PV expression in females compared to males (p < 0.0001, 95% C.I = [267746, 621366]; Figure 8D) and also reduced δ expression on PV neurons in females as compared to males (p < 0.0001, 95% C.I = [35568, 126727]; Figure 8E). These data demonstrate baseline sex differences in the expression and lability of GABA_A_R δ expression on PV neurons in the BLA.

**Figure 8.**
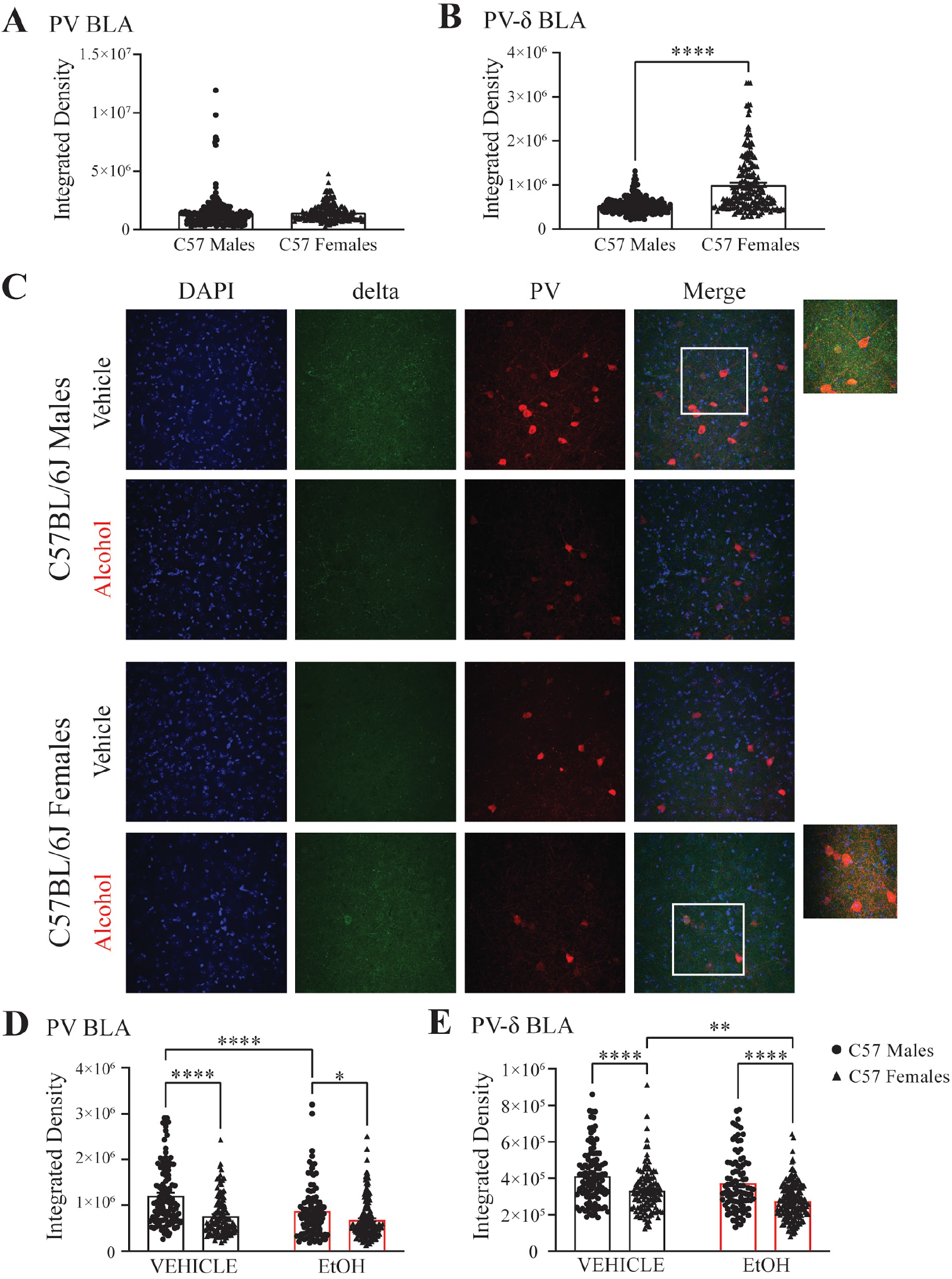
Repeated alcohol reduces δ-GABA_A_R expression on PV interneurons in female C57BL/6J mice. Integrated density of **A,** PV immunoreactivity and **B,** δ expression on PV interneurons in naive male (cell *n* = 224, animal *n* = 4) and female (cell *n* = 202, animal *n* = 4) C57BL/6J mice. **C,** Representative images from male and female C57BL/6J mice who received repeated vehicle or alcohol. Integrated density of **D,** PV immunoreactivity and **E,** δ expression on PV interneurons in the BLA of male (vehicle: cell *n* = 112, animal *n* = 3; alcohol: cell *n* = 97, animal *n* = 3) and female (vehicle: cell *n* = 117, animal *n* = 3; alcohol: cell *n* = 169, animal *n* = 3) C57BL/6J mice who received repeated vehicle or alcohol. *p < 0.05, **p < 0.01, ****p < 0.0001.

Repeated alcohol treatment reduced PV expression in males as compared to vehicle (p < 0.0001, 95% C.I = [147380, 518402]; Figure 8D). In contrast, repeated alcohol exposure in females did not alter PV expression but did significantly reduce GABA_A_R δ expression on PV interneurons compared to vehicle (p = 0.0018, 95% C.I = [15583, 98517]; Figure 8E), an effect that was not observed in males.

Comparing males and females exposed to repeated alcohol, PV expression (p = 0.0195, 95% C.I = [22163, 362911]; Figure 8D) and δ expression on PV interneurons are reduced in females as compared to males (p < 0.0001, 95% C.I = [54848, 142689; Figure 8E). These data implicate changes in GABA_A_R δ expression on PV interneurons in mediating sex differences and the response to repeated alcohol exposure.

## Discussion

Network states have been shown to correlate with behavioral states and accumulating evidence demonstrates that signature oscillatory states in the BLA are associated with fear and anxiety states (Likhtik et al., 2013; Stujenske et al., 2014; Davis et al., 2017; Antonoudiou et al., 2021). In fact, we demonstrated that optogenetically driving specific oscillatory states influences the behavioral expression of fear (Ozawa et al., 2020) and learned helplessness (Antonoudiou et al., 2021). However, limited studies have examined the physiological, pathological, or pharmacological mechanisms mediating transitions between network and behavioral states. Work from our laboratory has demonstrated that chronic stress can perturb oscillations in the BLA and a clinically effective antidepressant treatment can restore the “healthy” network state (Antonoudiou et al., 2021). Here, we examine the impact of alcohol on BLA network states. Given the anxiolytic effects of alcohol, we posited that alcohol may be capable of shifting the network state towards the anxiolytic state. We demonstrate that acute alcohol exposure is capable of altering BLA network states and that there are sex differences in the effect of alcohol on BLA network states, affecting different frequencies in males and females. These data are the first to demonstrate that alcohol is capable of modulating network states associated with affective states. Our lab has demonstrated that PV interneurons are critical in orchestrating oscillatory states in the BLA (Antonoudiou et al., 2021). PV interneurons in the BLA express a high density of δ-GABA_A_Rs, which have been suggested to be a target for low dose alcohol (Sundstrom-Poromaa et al., 2002; Wallner et al., 2003; Hanchar et al., 2006; Santhakumar et al., 2007). However, the actions of alcohol directly on these receptors remains somewhat controversial (Borghese et al., 2006; Korpi et al., 2007). It is important to note that the majority of these studies focus solely on principal neurons; GABAergic interneurons, on the other hand, have a unique receptor subunit composition in which the δ subunit has been shown to partner with the α1 subunit, and have been demonstrated to generate tonic GABAergic currents which are highly sensitive to low concentrations of ethanol (Glykys et al., 2007). Thus, we proposed that the high expression of δ subunit-containing GABA_A_Rs on PV interneurons in the BLA may confer unique sensitivity to the effects of alcohol and, given the role of these interneurons in coordinating oscillations, may mediate the effects of alcohol on BLA network states. Here we demonstrate that δ-GABA_A_Rs influence the ability of alcohol to alter specific oscillatory states in the BLA, blunting the ability to shift network states. Specifically, we observed a reduction of beta power from acute ethanol in male wild-type mice that was blunted in mice lacking δ-GABA_A_Rs (Figure 1, 3). This power detected in the beta frequency may arise from the neighboring high-theta oscillator, given the lack of a clear beta peak (Figure 1D). Regardless, these data suggest that δ subunit-containing GABA_A_Rs are important players in mediating the effects of alcohol on oscillatory states related to mood/anxiety; although, it is also possible that other GABA_A_R subtypes are involved. We previously demonstrated that δ has a specific role in lower frequencies as compared to higher frequencies (Antonoudiou et al., 2021), which may be true for the effects of alcohol as well. Further studies are required to investigate the impact of other GABA_A_R subtypes in mediating the ability of alcohol to modulate BLA network states given that previous studies have implicated other GABA_A_R subtypes, such as the γ2 subunit, in anxiety-like behavior (Chandra et al., 2005) and alcohol withdrawal severity (Buck and Hood, 1998). It is also possible that alcohol’s indirect effects on receptor expression, neurotransmitter availability, and other neuromodulators could account for the changes in BLA oscillations observed here (Morrow et al., 2001; Fleming et al., 2009; Olsen and Liang, 2017). For example, the effects of alcohol have been suggested to be mediated through the action of neuroactive steroids (Morrow et al., 2001; Finn et al., 2009; Finn and Jimenez, 2018) and given recent evidence that allopregnanolone can alter BLA network states (Antonoudiou et al., 2021), this may be an indirect mechanism whereby alcohol could modulate BLA network states. Arguing against this indirect mechanism is the evidence that alcohol exerts unique effects on BLA network states compared to allopregnanolone (Antonoudiou et al., 2021).

Since δ subunit-containing GABA_A_Rs have also been shown to be sensitive to ovarian steroid hormone modulation, we hypothesized that there may be sex differences in the ability of alcohol to modulate BLA network states through actions on these receptors. In fact, we do observe sex differences in the modulation of BLA network states by alcohol. Interestingly, the loss of the GABA_A_R δ subunit in males shifts the alcohol modulation of the BLA network state towards the signature that we observe for female C57BL/6J mice (Figure 4, 6) and we believe that the observed sex differences in the expression of δ subunit-containing GABA_A_Rs in the BLA (Figure 8) may underlie these differences. Further, there are well-documented sex differences in responses to alcohol, alcohol related anxiety-like behavior, and estrous-cycle dependent δ expression (Maguire et al., 2005; Rhodes et al., 2005; Barkley-Levenson and Crabbe, 2015), consistent with our observations of sex differences in the alcohol-induced modulation of network states. Additionally, sex differences have been reported in neural oscillations in major depressive disorder with oscillatory signatures of susceptibility (Thériault et al., 2021). Future studies are required to evaluate the relationship between the capacity of alcohol to modulate network states and voluntary alcohol consumption, the anxiolytic effects of alcohol, and the anxiogenic effects of alcohol withdrawal.

To investigate whether the effect of alcohol on network states may be altered after repeated exposure, we treated mice with low dose alcohol for up to five days. We found robust effects of repeated alcohol exposure by the last day of exposure on BLA LFP response of male C57BL/6J mice which was significantly different from the male *Gabrd^-/-^* mice and female C57BL/6J mice. In fact, we found no differences in the extent of the effect of repeated alcohol on network states in female C57BL/6J mice or *Gabrd^-/^* mice. Further, we found that BLA network states changed before alcohol exposure, suggesting some anticipation of the injections, which has been demonstrated in a Pavlovian conditioning paradigm (Tallot et al., 2020). Since the downregulation of the δ subunit has been thought to confer tolerance to alcohol (Olsen and Liang, 2017), the reduction of δ subunit in female, but not male, C57BL/6J mice could explain the lack of LFP effect after repeated alcohol exposure and potentially point to a mechanism contributing to tolerance in females. It is also possible that δ expression in female C57BL/6J mice is so low that alcohol’s effects through δ-GABA_A_Rs are too small to affect BLA oscillations, which could explain why the female C57BL/6J effects are similar to *Gabrd^-/^* mice. Further, we found that δ expression on PV interneurons is increased in naive female C57BL/6J mice compared to males. Because we did not see any baseline differences in BLA network states between male and female C57BL/6J mice, this difference in expression may not impact BLA oscillations, but expression of δ in females does influence the response to alcohol exposure.

The literature and our recent findings demonstrate a strong role for PV interneurons in oscillation generation (Antonoudiou et al., 2021) giving support to the likely fact that alcohol’s effects on PV interneurons are influencing the oscillations. However, due to the heterogeneity of the interneuron population in the BLA, it is possible other interneuron types, like somatostatin, cholecystokinin, or PKC-δ expressing cells may be involved in effecting oscillations (Klausberger et al., 2005). Furthermore, another major influence on BLA oscillations are other brain areas with strong network connections to the BLA, such as the medial prefrontal cortex (mPFC) (Davis et al., 2017; Ozawa et al., 2020).

Forced alcohol injections or alcohol induced aversion can cause stress to mice which may contribute to the observed effects (Eckardt et al., 1974). However, our experimental plan was designed to dissociate any stressful or unpleasant effects of the infusion from the real effects of alcohol. We did observe significant effects of acute vehicle injections on BLA LFP responses as compared to baseline (Figure 1C, 2A, 3A), but did not find any sensitization or tolerance to the injections in the acute alcohol experiment or across days in the repeated alcohol experiment (Figure 2-1, 3-1, 4-1). This is similar to what has been reported previously from our lab (Antonoudiou et al., 2021). Thus, we are confident that the observed effects of alcohol on BLA oscillatory states is due to the effects of alcohol rather than an aversive experience due to the route of administration. Further, our data suggest that the effects of alcohol may mitigate the stress-induced effects on the BLA network state.

To our knowledge, this is the first demonstration that alcohol can modulate oscillations in the BLA, which have been implicated in governing behavioral states. Numerous studies have investigated the relationship of BLA network states to behavioral states; however, few studies have investigated mechanisms mediating transitions between BLA network states. The current study demonstrates that alcohol can induce a transition between network states associated with fear and anxiety, which may mediate the impact of alcohol on anxiety states. Future work is required to investigate how changes in the BLA relate to other connected areas implicated in alcohol use and anxiety, such as the central amygdala, mPFC, nucleus accumbens, BNST, and ventral striatum (Janak and Tye, 2015). Recordings of oscillations are stable over long periods of time and thus can be examined throughout the addiction cycle from intoxication to withdrawal to preoccupation in specific brain areas to understand how alcohol changes communication between these areas. Thus, this novel approach may demonstrate utility in understanding the trajectory from first drink to alcohol dependence and the contribution of both the positive and negative reinforcing effects of alcohol.

## Acknowledgements

The authors would like to thank Dr. Klaus Miczek, Dr. Laverne Melón, and Dr. Leon Reijmers for thoughtful feedback and guidance on this project. Visual abstract created with BioRender.com.

