## Supplemental Materials for "Sex differences in the alcohol-mediated modulation of BLA network states"

**Extended Data Figures & Tables**
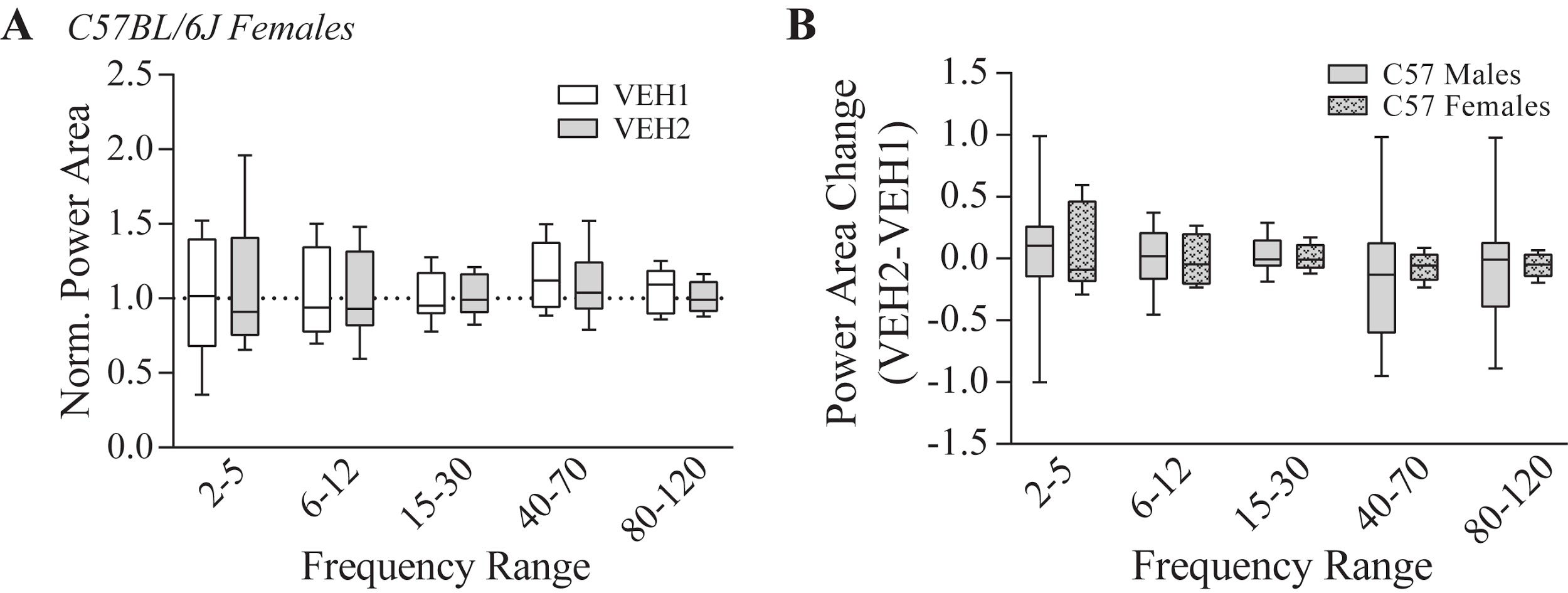

**Figure 2-1.** Acute vehicle injections do not alter BLA LFP networks in female C57BL/6J mice. **A,** Normalized power area for acute vehicle/vehicle exposure in female C57BL/6J mice (*n* = 6). **B,** Power area difference between the first and second vehicle injection in male and female C57BL/6J mice.

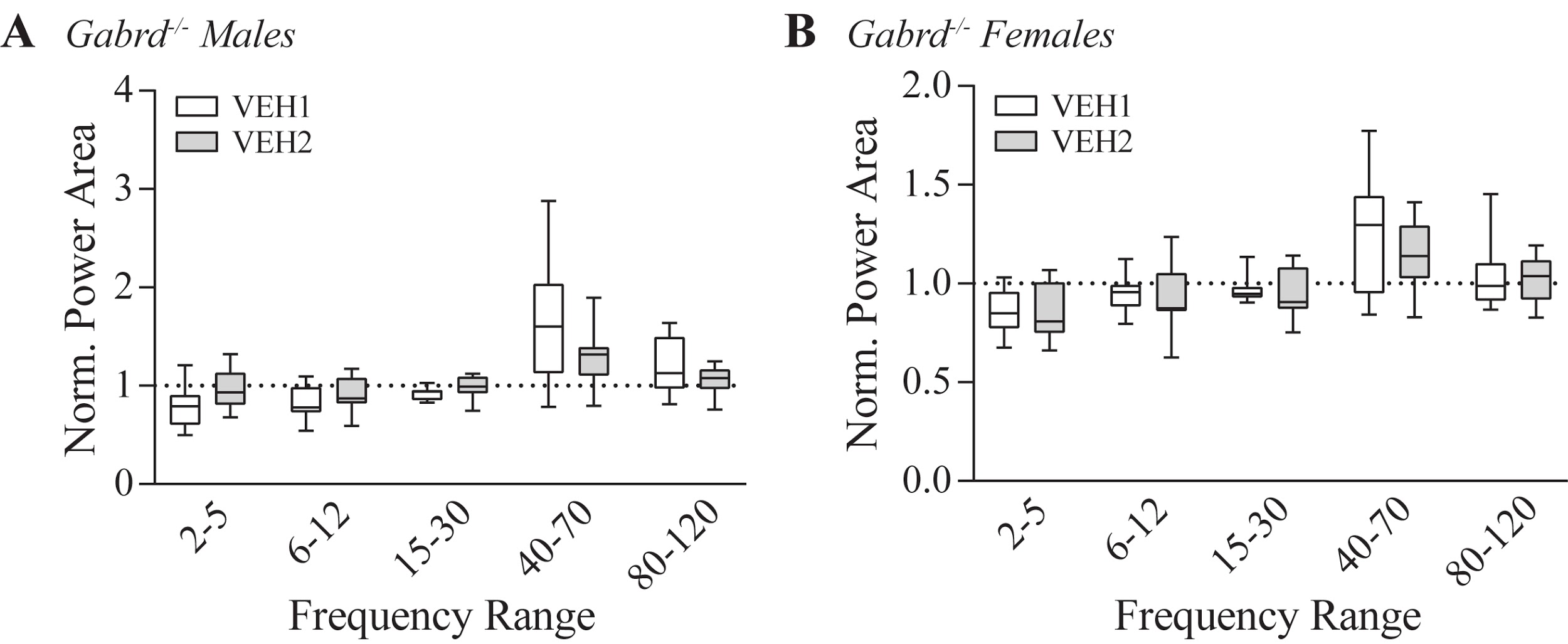

**Figure 3-1.** Vehicle injections do not alter BLA LFP networks in Gabrd^-/-^ mice. **A,** Normalized power area for vehicle/vehicle exposure in male Gabrd^-/-^ mice (*n* = 8) and **B,** female Gabrd^-/-^ mice (*n* = 7).

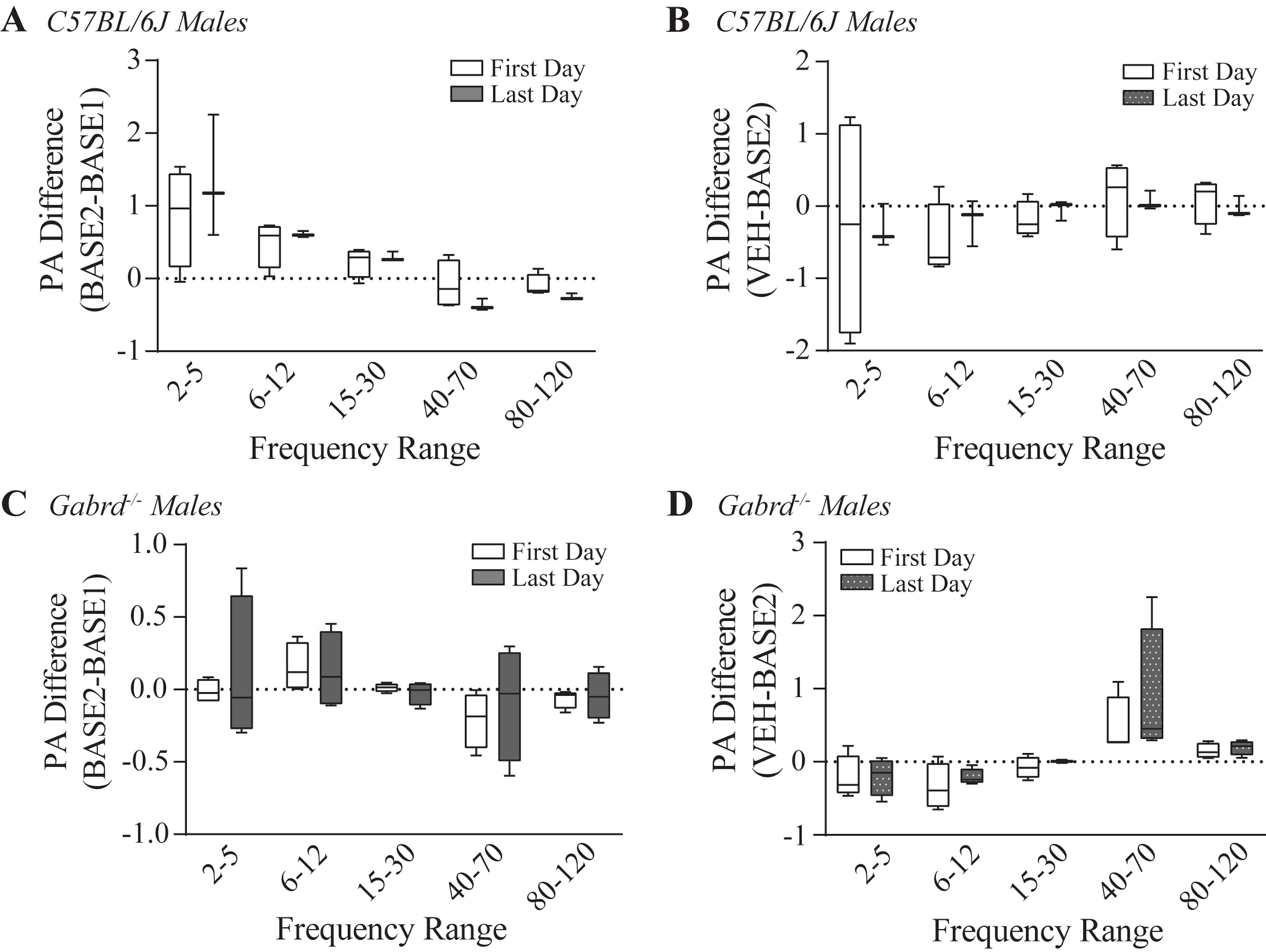

**Figure 4-1.** Repeated vehicle injections do not change BLA LFP network states in male C57BL6/J and Gabrd^-/-^ mice. Power area difference within baseline for first and last day of exposure in **A,** male C57BL/6J (first day *n* = 4; last day *n* = 3) and **C,** male Gabrd^-/-^mice (first day *n* = 4; last day *n* = 4). Power area difference between vehicle and baseline on the first and last day of exposure in **B,** male C57BL/6J mice and **D,** male Gabrd^-/-^ mice.

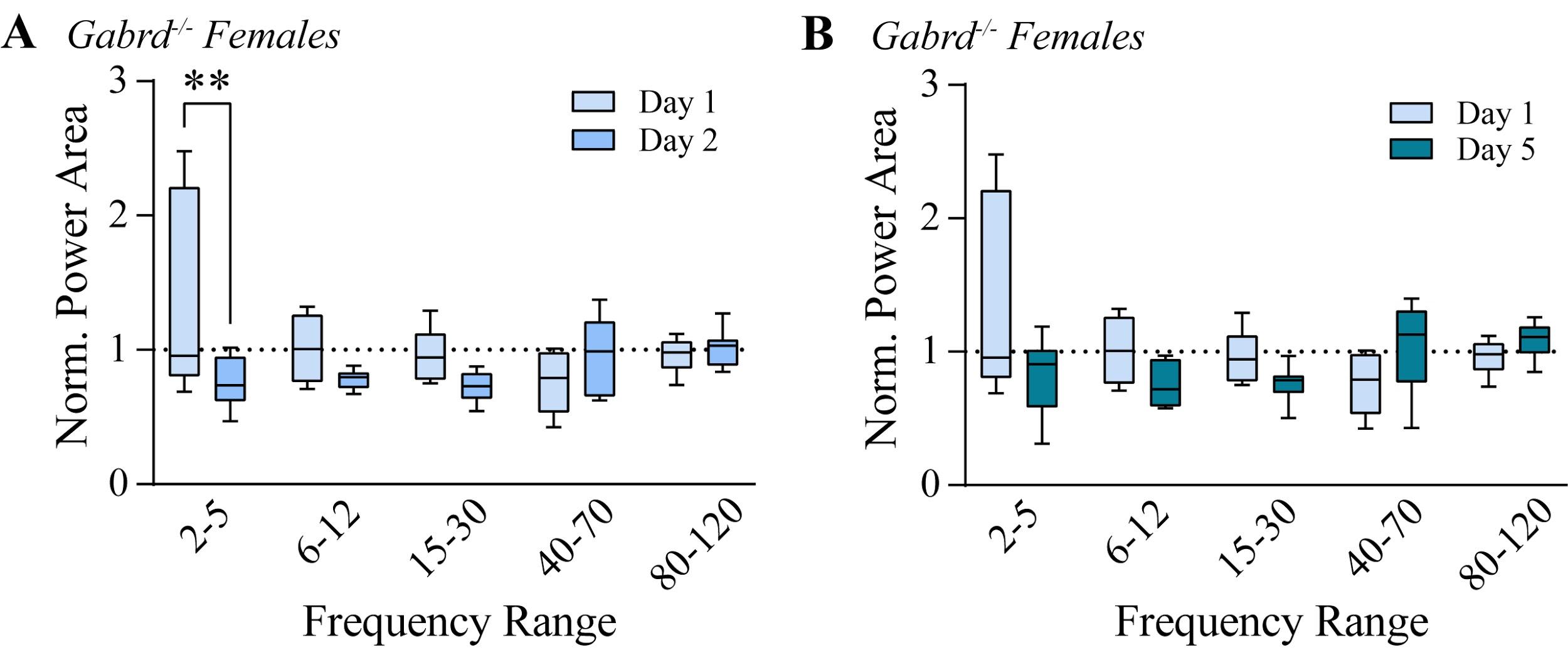
**Figure 6-1.** Effects of alcohol on day one (acute) and day five do not differ in female Gabrd^-/-^ mice. **A,** Normalized power area of BLA LFP of alcohol injections during the acute alcohol injection (day one) and day two in female Gabrd^-/-^ mice. **B,** Normalized power area of BLA LFP of alcohol injections during acute alcohol injection (day one) and day five of repeated alcohol injections in female Gabrd^-/-^ mice. **p < 0.01.

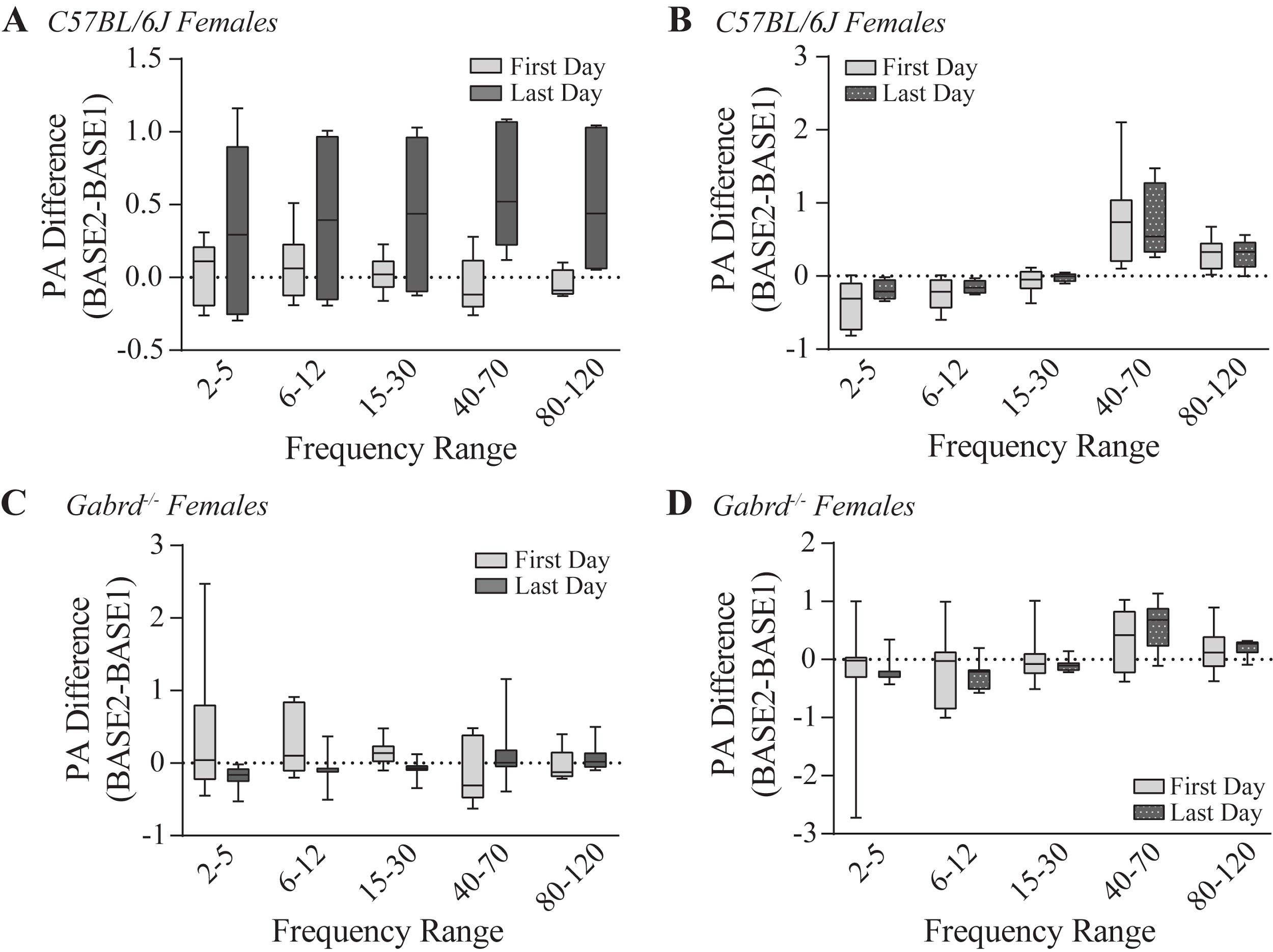

**Figure 6-2.** Repeated vehicle injections do not change BLA LFP networks over days in female C57BL/6J and Gabrd^-/-^ mice. Power area difference within baseline from the first to the last day of exposure in **A,** female C57BL/6J (first day *n* = 5; last day *n* = 6) and **C,** female Gabrd^-/-^ mice (first day *n* = 7; last day *n* = 7). Power area difference between vehicle and baseline from the first to the last day of exposure in **B,** female C57BL/6J and **D,** female Gabrd^-/-^ mice.

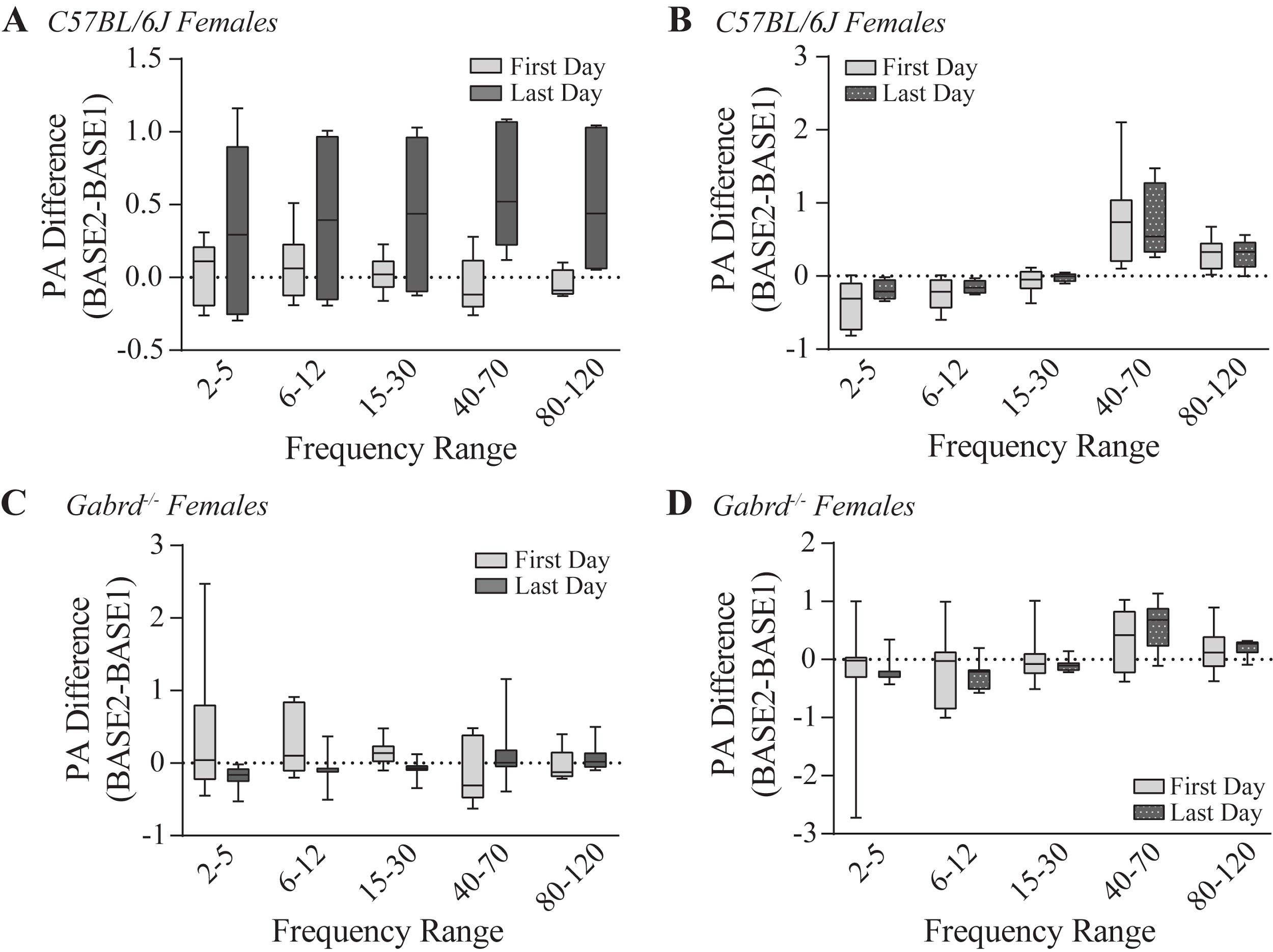

**Figure 6-3.** No differences between effects of repeated alcohol on BLA LFP network states in female C57BL/6J or Gabrd^-/-^ mice. **A,** Change in baseline on the first and **C,** last day of exposure between female C57BL/6J and Gabrd^-/-^ mice. **B,** Change in effect of alcohol on the first and **D,** last day of exposure between female C57BL/6J and Gabrd^-/-^ mice.

**Table 1-1.** Summary of ANOVA tests for acute alcohol experiments.

| **Description** | **Figure** | **Factors** | **SS** | **DF** | **MS** | **F (DFn, DFd)** | **P value** |
| --- | --- | --- | --- | --- | --- | --- | --- |
| Acute VEH: C57BL/6J male (veh vs veh) | Fig. 1C | Frequency | 11.48 | 4 | 2.87 | F (1.551, 18.61) = 19.16 | 0.0001 |
|  |  | Treatment | 0.0434 | 1 | 0.0434 | F (1.000, 12.00) = 0.5896 | 0.4574 |
|  |  | Frequency x Treatment | 0.1914 | 4 | 0.04784 | F (1.144, 13.73) = 0.5730 | 0.4842 |
| Acute VEH: C57BL/6J male (veh vs baseline) | Fig. 1C | Frequency | 3.64 | 4 | 0.9099 | F (1.645, 19.74) = 19.94 | 0.0001 |
|  |  | Treatment | 0.2267 | 1 | 0.2267 | F (1.000, 12.00) = 3.690 | 0.0788 |
|  |  | Frequency x Treatment | 3.64 | 4 | 0.9099 | F (1.645, 19.74) = 19.94 | 0.0001 |
| Acute EtOH: C57BL/6J male (veh vs etoh) | Fig. 1E | Frequency | 3.858 | 4 | 0.9644 | F (1.137, 11.37) = 2.913 | 0.1122 |
|  |  | Treatment | 0.6475 | 1 | 0.6475 | F (1.000, 10.00) = 13.37 | 0.0044 |
|  |  | Frequency x Treatment | 0.01328 | 4 | 0.00332 | F (1.237, 12.37) = 0.07865 | 0.8345 |
| Acute VEH: C57BL/6J female (veh vs veh) | Ext. Fig. 2-1A | Frequency | 0.08763 | 4 | 0.02191 | F (1.269, 6.345) = 0.1491 | 0.7695 |
|  |  | Treatment | 0.002078 | 1 | 0.00208 | F (1.000, 5.000) = 0.08908 | 0.7774 |
|  |  | Frequency x Treatment | 0.03402 | 4 | 0.00851 | F (1.097, 5.484) = 0.4288 | 0.5572 |
| Acute VEH: C57BL/6J female (veh vs baseline) | Ext. Fig. 2-1A | Frequency | 0.0443 | 4 | 0.01107 | F (1.198, 5.990) = 0.2681 | 0.6643 |
|  |  | Treatment | 0.0391 | 1 | 0.0391 | F (1.000, 5.000) = 1.264 | 0.312 |
|  |  | Frequency x Treatment | 0.0443 | 4 | 0.01107 | F (1.198, 5.990) = 0.2681 | 0.6643 |
| Acute EtOH: C57BL/6J female (veh vs etoh) | Fig. 2A | Frequency | 3.145 | 4 | 0.7861 | F (1.153, 8.069) = 7.026 | 0.0263 |
|  |  | Treatment | 0.4585 | 1 | 0.4585 | F (1.000, 7.000) = 18.34 | 0.0036 |
|  |  | Frequency x Treatment | 1.307 | 4 | 0.3268 | F (1.669, 11.68) = 12.78 | 0.0016 |
| Acute EtOH: C57BL/6J female (veh vs baseline) | Fig. 2A | Frequency | 1.981 | 4 | 0.4952 | F (1.237, 8.660) = 10.86 | 0.0077 |
|  |  | Treatment | 0.1065 | 1 | 0.1065 | F (1.000, 7.000) = 6.002 | 0.0441 |
|  |  | Frequency x Treatment | 1.981 | 4 | 0.4952 | F (1.237, 8.660) = 10.86 | 0.0077 |
| Acute VEH: C57BL/6J male vs female | Ext. Fig. 2-1B | Frequency x Sex | 0.04076 | 4 | 0.01019 | F (4, 68) = 0.07867 | 0.9886 |
|  |  | Frequency | 0.2941 | 4 | 0.07352 | F (1.145, 19.47) = 0.5676 | 0.4828 |
|  |  | Sex | 0.0126 | 1 | 0.0126 | F (1, 17) = 0.1071 | 0.7475 |
|  |  | Subject | 2 | 17 | 0.1176 | F (17, 68) = 0.9081 | 0.5675 |
| Acute EtOH: C57BL/6J male vs female | Fig. 2B | Frequency x Sex | 1.418 | 4 | 0.3544 | F (4, 80) = 3.479 | 0.0113 |
|  |  | Frequency | 2.014 | 4 | 0.5036 | F (1.453, 29.07) = 4.943 | 0.0225 |
|  |  | Sex | 0.07188 | 1 | 0.07188 | F (1, 20) = 0.4966 | 0.4891 |
|  |  | Subject | 2.895 | 20 | 0.1448 | F (20, 80) = 1.421 | 0.1373 |
| Acute VEH: Gabrd^-/-^ male (veh vs veh) | Ext. Fig. 3-1A | Frequency | 4.086 | 4 | 1.021 | F (1.641, 11.49) = 8.285 | 0.0078 |
|  |  | Treatment | 0.02705 | 1 | 0.02705 | F (1.000, 7.000) = 1.526 | 0.2565 |
|  |  | Frequency x Treatment | 0.6961 | 4 | 0.174 | F (1.327, 9.287) = 4.270 | 0.0601 |
| Acute VEH: Gabrd^-/-^ male (veh vs baseline) | Ext. Fig. 3-1A | Frequency | 2.018 | 4 | 0.5044 | F (1.570, 10.99) = 8.012 | 0.0097 |
|  |  | Treatment | 0.1178 | 1 | 0.1178 | F (1.000, 7.000) = 2.334 | 0.1705 |
|  |  | Frequency x Treatment | 2.018 | 4 | 0.5044 | F (1.570, 10.99) = 8.012 | 0.0097 |
| Acute EtOH: Gabrd^-/-^ male (veh vs etoh) | Fig. 3A | Frequency | 5.668 | 4 | 1.417 | F (1.452, 11.62) = 11.46 | 0.0031 |
|  |  | Treatment | 0.7003 | 1 | 0.7003 | F (1.000, 8.000) = 138.9 | 0.0001 |
|  |  | Frequency x Treatment | 0.4209 | 4 | 0.1052 | F (1.190, 9.519) = 2.631 | 0.1351 |
| Acute EtOH: Gabrd^-/-^ male (vs baseline) | Fig. 3A | Frequency | 2.24 | 4 | 0.5599 | F (1.331, 10.64) = 9.112 | 0.0085 |
|  |  | Treatment | 0.2227 | 1 | 0.2227 | F (1.000, 8.000) = 11.94 | 0.0086 |
|  |  | Frequency x Treatment | 2.24 | 4 | 0.5599 | F (1.331, 10.64) = 9.112 | 0.0085 |
| Acute VEH: Gabrd^-/-^ female (veh vs veh) | Ext. Fig. 3-1B | Frequency | 0.9047 | 4 | 0.2262 | F (1.171, 7.025) = 5.615 | 0.0461 |
|  |  | Treatment | 0.02948 | 1 | 0.02948 | F (1.000, 6.000) = 6.270 | 0.0463 |
|  |  | Frequency x Treatment | 0.02111 | 4 | 0.00528 | F (1.143, 6.856) = 0.3050 | 0.628 |
| Acute VEH: Gabrd^-/-^ female (veh vs baseline) | Ext. Fig. 3-1B | Frequency | 0.2981 | 4 | 0.07452 | F (1.185, 7.108) = 4.097 | 0.0783 |
|  |  | Treatment | 0.004764 | 1 | 0.00476 | F (1.000, 6.000) = 0.4050 | 0.548 |
|  |  | Frequency x Treatment | 0.2981 | 4 | 0.07452 | F (1.185, 7.108) = 4.097 | 0.0783 |
| Acute EtOH: Gabrd^-/-^ female (veh vs etoh) | Fig. 3B | Frequency | 2.293 | 4 | 0.5733 | F (1.056, 7.392) = 1.779 | 0.2235 |
|  |  | Treatment | 0.4624 | 1 | 0.4624 | F (1.000, 7.000) = 11.31 | 0.012 |
|  |  | Frequency x Treatment | 0.2087 | 4 | 0.05217 | F (1.289, 9.023) = 2.638 | 0.1353 |
| Acute EtOH: Gabrd^-/-^ female (veh vs baseline) | Fig. 3B | Frequency | 0.4223 | 4 | 0.1056 | F (1.064, 7.449) = 1.073 | 0.3383 |
|  |  | Treatment | 0.5712 | 1 | 0.5712 | F (1.000, 7.000) = 4.697 | 0.0669 |
|  |  | Frequency x Treatment | 0.4223 | 4 | 0.1056 | F (1.064, 7.449) = 1.073 | 0.3383 |
| Acute EtOH: C57BL/6J male vs Gabrd^-/-^ male | Fig. 3C | Frequency x Genotype | 0.7897 | 4 | 0.1974 | F (4, 76) = 1.933 | 0.1135 |
|  |  | Frequency | 0.3485 | 4 | 0.08712 | F (1.362, 25.89) = 0.8532 | 0.3978 |
|  |  | Genotype | 0.007213 | 1 | 0.00721 | F (1, 19) = 0.06556 | 0.8007 |
|  |  | Subject | 2.091 | 19 | 0.11 | F (19, 76) = 1.078 | 0.39 |
| Acute EtOH: C57BL/6J female vs Gabrd^-/-^ female | Fig. 3D | Frequency x Genotype | 0.5119 | 4 | 0.128 | F (4, 56) = 2.822 | 0.0333 |
|  |  | Frequency | 2.52 | 4 | 0.63 | F (1.596, 22.35) = 13.89 | 0.0003 |
|  |  | Genotype | 8.11E-06 | 1 | 8.1E-06 | F (1, 14) = 0.0001230 | 0.9913 |
|  |  | Subject | 0.9224 | 14 | 0.06588 | F (14, 56) = 1.453 | 0.1603 |

**Table 1-2.** Summary of Šídák's multiple comparisons tests for acute alcohol experiments.

| **Description** | **Figure** | **Comparisons** | **P value** | **Mean Diff** | **SEM** | ***n*** |
| --- | --- | --- | --- | --- | --- | --- |
| Acute VEH: C57BL/6J male (veh vs veh) | Fig. 1C | 2-5 | >0.9999 | -0.02528 | 0.1347 | 13 |
|  |  | 6-12 | 0.9997 | -0.01681 | 0.06667 | 13 |
|  |  | 15-30 | 0.9545 | -0.03081 | 0.04046 | 13 |
|  |  | 40-70 | 0.8412 | 0.1559 | 0.1464 | 13 |
|  |  | 80-120 | 0.9539 | 0.09971 | 0.1305 | 13 |
| Acute VEH: C57BL/6J male (veh vs baseline) | Fig. 1C | 2-5 | 0.1323 | 0.2264 | 0.0906 | 13 |
|  |  | 6-12 | 0.0117 | 0.2248 | 0.05854 | 13 |
|  |  | 15-30 | 0.082 | 0.0894 | 0.03228 | 13 |
|  |  | 40-70 | 0.0027 | -0.5932 | 0.127 | 13 |
|  |  | 80-120 | 0.0103 | -0.3649 | 0.09336 | 13 |
| Acute EtOH: C57BL/6J male (veh vs etoh) | Fig. 1E | 2-5 | 0.7302 | 0.1503 | 0.1177 | 11 |
|  |  | 6-12 | 0.2567 | 0.1711 | 0.07979 | 11 |
|  |  | 15-30 | 0.034 | 0.1514 | 0.04469 | 11 |
|  |  | 40-70 | 0.3611 | 0.1789 | 0.09386 | 11 |
|  |  | 80-120 | 0.7432 | 0.1155 | 0.09206 | 11 |
| Acute VEH: C57BL/6J female (veh vs veh) | Ext. Fig. 2-1A | 2-5 | 0.9962 | -0.06576 | 0.1461 | 6 |
|  |  | 6-12 | >0.9999 | 0.01346 | 0.0831 | 6 |
|  |  | 15-30 | 0.9998 | -0.01017 | 0.04417 | 6 |
|  |  | 40-70 | 0.7365 | 0.06639 | 0.04908 | 6 |
|  |  | 80-120 | 0.7317 | 0.05493 | 0.04033 | 6 |
| Acute VEH: C57BL/6J female (veh vs baseline) | Fig. 2A | 2-5 | >0.9999 | -0.007645 | 0.1735 | 6 |
|  |  | 6-12 | 0.9999 | -0.02814 | 0.1266 | 6 |
|  |  | 15-30 | >0.9999 | -0.005704 | 0.07185 | 6 |
|  |  | 40-70 | 0.6555 | -0.152 | 0.1008 | 6 |
|  |  | 80-120 | 0.9006 | -0.06182 | 0.06276 | 6 |
| Acute EtOH: C57BL/6J female (veh vs etoh) | Fig. 2A | 2-5 | 0.8009 | -0.1429 | 0.1209 | 8 |
|  |  | 6-12 | >0.9999 | 0.01077 | 0.05909 | 8 |
|  |  | 15-30 | 0.2422 | 0.1183 | 0.05114 | 8 |
|  |  | 40-70 | 0.0014 | 0.6198 | 0.09307 | 8 |
|  |  | 80-120 | 0.0883 | 0.1511 | 0.04938 | 8 |
| Acute EtOH: C57BL/6J female (veh vs baseline) | Fig. 2A | 2-5 | 0.1081 | 0.2115 | 0.07266 | 8 |
|  |  | 6-12 | 0.0273 | 0.2283 | 0.05777 | 8 |
|  |  | 15-30 | 0.6743 | 0.03529 | 0.025 | 8 |
|  |  | 40-70 | 0.0557 | -0.6083 | 0.1787 | 8 |
|  |  | 80-120 | 0.1985 | -0.2316 | 0.09406 | 8 |
| Acute VEH: C57BL/6J male vs female | Ext. Fig. 2-1B | 2-5 | >0.9999 | -0.04048 | 0.1987 | 13, 6 |
|  |  | 6-12 | 0.9995 | 0.03027 | 0.1065 | 13, 6 |
|  |  | 15-30 | 0.9987 | 0.02064 | 0.05989 | 13, 6 |
|  |  | 40-70 | 0.9855 | -0.08951 | 0.1544 | 13, 6 |
|  |  | 80-120 | 0.999 | -0.04479 | 0.1366 | 13, 6 |
| Acute EtOH: C57BL/6J male vs female | Fig. 2B | 2-5 | 0.3389 | -0.3646 | 0.1973 | 14, 8 |
|  |  | 6-12 | 0.2586 | -0.1988 | 0.09891 | 14, 8 |
|  |  | 15-30 | 0.9551 | -0.04979 | 0.06609 | 14, 8 |
|  |  | 40-70 | 0.1425 | 0.343 | 0.1469 | 14, 8 |
|  |  | 80-120 | >0.9999 | 0.004589 | 0.08976 | 14, 8 |
| Acute VEH: Gabrd^-/-^ male (veh vs veh) | Ext. Fig. 3-1A | 2-5 | 0.5861 | -0.1647 | 0.1053 | 8 |
|  |  | 6-12 | 0.8202 | -0.0894 | 0.0782 | 8 |
|  |  | 15-30 | 0.5514 | -0.06128 | 0.03771 | 8 |
|  |  | 40-70 | 0.2407 | 0.3426 | 0.1479 | 8 |
|  |  | 80-120 | 0.2522 | 0.1566 | 0.06862 | 8 |
| Acute VEH: Gabrd^-/-^ male (veh vs baseline) | Ext. Fig. 3-1A | 2-5 | 0.1649 | 0.2036 | 0.0783 | 8 |
|  |  | 6-12 | 0.1067 | 0.1828 | 0.06258 | 8 |
|  |  | 15-30 | 0.0585 | 0.07961 | 0.02366 | 8 |
|  |  | 40-70 | 0.1352 | -0.6389 | 0.2326 | 8 |
|  |  | 80-120 | 0.3513 | -0.2107 | 0.1042 | 8 |
| Acute EtOH: Gabrd^-/-^ male (veh vs etoh) | Fig. 3A | 2-5 | 0.9997 | 0.02016 | 0.07707 | 9 |
|  |  | 6-12 | 0.1498 | 0.1159 | 0.04469 | 9 |
|  |  | 15-30 | 0.2742 | 0.1149 | 0.05304 | 9 |
|  |  | 40-70 | 0.0202 | 0.4227 | 0.1063 | 9 |
|  |  | 80-120 | 0.4815 | 0.2084 | 0.1209 | 9 |
| Acute EtOH: Gabrd^-/-^ male (veh vs baseline) | Fig. 3A | 2-5 | 0.2764 | 0.2146 | 0.0993 | 9 |
|  |  | 6-12 | 0.7899 | 0.1183 | 0.09939 | 9 |
|  |  | 15-30 | 0.6008 | 0.1085 | 0.07155 | 9 |
|  |  | 40-70 | 0.0121 | -0.601 | 0.138 | 9 |
|  |  | 80-120 | 0.1156 | -0.3379 | 0.1219 | 9 |
| Acute VEH: Gabrd^-/-^ female (veh vs veh) | Ext. Fig. 3-1B | 2-5 | >0.9999 | 0.008554 | 0.08033 | 7 |
|  |  | 6-12 | 0.9898 | 0.02593 | 0.04695 | 7 |
|  |  | 15-30 | 0.9753 | 0.02565 | 0.03781 | 7 |
|  |  | 40-70 | 0.8039 | 0.1081 | 0.09063 | 7 |
|  |  | 80-120 | 0.9723 | 0.03701 | 0.05312 | 7 |
| Acute VEH: Gabrd^-/-^ female (veh vs baseline) | xt. Fig. 3-1B | 2-5 | 0.1004 | 0.1395 | 0.04492 | 7 |
|  |  | 6-12 | 0.7362 | 0.05026 | 0.03798 | 7 |
|  |  | 15-30 | 0.8999 | 0.02779 | 0.02861 | 7 |
|  |  | 40-70 | 0.3637 | -0.2468 | 0.1205 | 7 |
|  |  | 80-120 | 0.9668 | -0.05326 | 0.07312 | 7 |
| Acute EtOH: Gabrd^-/-^ female (veh vs etoh) | Fig. 3B | 2-5 | 0.9854 | 0.07624 | 0.1282 | 8 |
|  |  | 6-12 | 0.6618 | 0.09839 | 0.06865 | 8 |
|  |  | 15-30 | 0.3425 | 0.1218 | 0.05963 | 8 |
|  |  | 40-70 | 0.0012 | 0.3541 | 0.05196 | 8 |
|  |  | 80-120 | 0.2988 | 0.1097 | 0.051 | 8 |
| Acute EtOH: Gabrd^-/-^ female (veh vs baseline) | Fig. 3B | 2-5 | 0.6359 | -0.4575 | 0.3095 | 8 |
|  |  | 6-12 | 0.8335 | -0.1201 | 0.1077 | 8 |
|  |  | 15-30 | 0.8451 | -0.09036 | 0.08281 | 8 |
|  |  | 40-70 | 0.8855 | -0.109 | 0.1093 | 8 |
|  |  | 80-120 | 0.7168 | -0.06802 | 0.05087 | 8 |
| Acute EtOH: C57BL/6J male vs Gabrd^-/-^ male | Fig. 3C | 2-5 | 0.6317 | -0.2673 | 0.1906 | 12, 9 |
|  |  | 6-12 | 0.8322 | -0.107 | 0.09995 | 12, 9 |
|  |  | 15-30 | 0.96 | -0.04991 | 0.06825 | 12, 9 |
|  |  | 40-70 | 0.3741 | 0.2462 | 0.1365 | 12, 9 |
|  |  | 80-120 | 0.9774 | 0.09433 | 0.1473 | 12, 9 |
| Acute EtOH: C57BL/6J female vs Gabrd^-/-^ female | Fig. 3D | 2-5 | 0.7365 | 0.2191 | 0.1762 | 8, 8 |
|  |  | 6-12 | 0.8841 | 0.08761 | 0.09058 | 8, 8 |
|  |  | 15-30 | >0.9999 | 0.003523 | 0.07856 | 8, 8 |
|  |  | 40-70 | 0.1411 | -0.2657 | 0.1066 | 8, 8 |
|  |  | 80-120 | 0.9851 | -0.0414 | 0.07098 | 8, 8 |

**Table 4-1.** Summary of ANOVA tests for repeated alcohol experiments.

| **Description** | **Figure** | **Factors** | **SS** | **DF** | **MS** | **F (DFn, DFd)** | **P value** |
| --- | --- | --- | --- | --- | --- | --- | --- |
| Repeated EtOH: C57BL/6J male (Day 1 vs Day 5 - within base) | Figure 4C | Frequency x Day | 1.671 | 4 | 0.4176 | F (4, 48) = 1.915 | 0.1232 |
|  |  | Frequency | 8.35 | 4 | 2.087 | F (1.244, 14.93) = 9.572 | 0.0052 |
|  |  | Day | 0.1573 | 1 | 0.1573 | F (1, 12) = 0.4664 | 0.5076 |
|  |  | Subject | 4.047 | 12 | 0.3372 | F (12, 48) = 1.546 | 0.1407 |
| Repeated EtOH: C57BL/6J male (Day 1 vs Day 5 - from base) | Figure 4D | Frequency x Day | 3.175 | 4 | 0.7938 | F (4, 48) = 4.138 | 0.0058 |
|  |  | Frequency | 8.473 | 4 | 2.118 | F (1.178, 14.13) = 11.04 | 0.0037 |
|  |  | Day | 0.4699 | 1 | 0.4699 | F (1, 12) = 1.225 | 0.2901 |
|  |  | Subject | 4.604 | 12 | 0.3837 | F (12, 48) = 2.000 | 0.0451 |
| Repeated EtOH: Gabrd^-/-^ male (Day 1 vs Day 5 - within base) | Figure 4E | Frequency | 0.04903 | 4 | 0.01226 | F (1.230, 8.610) = 0.1890 | 0.7246 |
|  |  | Day | 0.07789 | 1 | 0.07789 | F (1.000, 7.000) = 1.650 | 0.2398 |
|  |  | Frequency x Day | 0.3835 | 4 | 0.09588 | F (1.204, 8.426) = 1.140 | 0.3297 |
| Repeated EtOH: Gabrd^-/-^ male (Day 1 vs Day 5 - from base) | Figure 4F | Frequency | 4.022 | 4 | 1.005 | F (1.340, 9.383) = 13.28 | 0.0033 |
|  |  | Day | 0.00048 | 1 | 0.00048 | F (1.000, 7.000) = 0.01124 | 0.9185 |
|  |  | Frequency x Day | 0.8977 | 4 | 0.2244 | F (1.322, 9.256) = 6.531 | 0.0244 |
| Repeated VEH: C57BL/6J male (Day 1 vs Day 5 - within base) | Ext. Fig. 4-1A | Frequency x Day | 0.6028 | 4 | 0.1507 | F (4, 20) = 1.014 | 0.4237 |
|  |  | Frequency | 8.249 | 4 | 2.062 | F (1.298, 6.490) = 13.88 | 0.0067 |
|  |  | Day | 0.02047 | 1 | 0.02047 | F (1, 5) = 0.1456 | 0.7184 |
|  |  | Subject | 0.7029 | 5 | 0.1406 | F (5, 20) = 0.9462 | 0.4731 |
| Repeated VEH: C57BL/6J male (Day 1 vs Day 5 - from base) | Ext. Fig. 4-1B | Frequency x Day | 0.1939 | 4 | 0.04848 | F (4, 20) = 0.1269 | 0.971 |
|  |  | Frequency | 1.043 | 4 | 0.2606 | F (1.050, 5.249) = 0.6820 | 0.4521 |
|  |  | Day | 0.02203 | 1 | 0.02203 | F (1, 5) = 0.04891 | 0.8337 |
|  |  | Subject | 2.252 | 5 | 0.4504 | F (5, 20) = 1.178 | 0.3544 |
| Repeated VEH: Gabrd^-/-^ male (Day 1 vs Day 5 - within base) | Ext. Fig. 4-1C | Frequency | 0.3759 | 4 | 0.09399 | F (1.037, 3.111) = 0.8809 | 0.4196 |
|  |  | Day | 0.01559 | 1 | 0.01559 | F (1.000, 3.000) = 1.315 | 0.3346 |
|  |  | Frequency x Day | 0.04469 | 4 | 0.01117 | F (1.168, 3.504) = 0.3053 | 0.6475 |
| Repeated VEH: Gabrd^-/-^ male (Day 1 vs Day 5 - from base) | Ext. Fig. 4-1D | Frequency | 4.601 | 4 | 1.15 | F (1.351, 4.053) = 18.62 | 0.0106 |
|  |  | Day | 0.1827 | 1 | 0.1827 | F (1.000, 3.000) = 3.547 | 0.1562 |
|  |  | Frequency x Day | 0.178 | 4 | 0.0445 | F (1.051, 3.152) = 0.2310 | 0.6735 |
| Repeated EtOH: C57BL/6J male vs Gabrd^-/-^ male (Day 1 - within base) | Figure 5A | Frequency x Genotype | 0.3485 | 4 | 0.08714 | F (4, 56) = 0.7633 | 0.5536 |
|  |  | Frequency | 1.575 | 4 | 0.3937 | F (1.361, 19.05) = 3.449 | 0.0679 |
|  |  | Genotype | 0.06296 | 1 | 0.06296 | F (1, 14) = 0.5065 | 0.4883 |
|  |  | Subject | 1.74 | 14 | 0.1243 | F (14, 56) = 1.089 | 0.3873 |
| Repeated EtOH: C57BL/6J male vs Gabrd^-/-^ male (Day 1 - from base) | Figure 5B | Frequency x Genotype | 1.158 | 4 | 0.2896 | F (4, 56) = 2.405 | 0.0603 |
|  |  | Frequency | 0.2006 | 4 | 0.05015 | F (1.203, 16.84) = 0.4166 | 0.5644 |
|  |  | Genotype | 1.74 | 1 | 1.74 | F (1, 14) = 9.046 | 0.0094 |
|  |  | Subject | 2.692 | 14 | 0.1923 | F (14, 56) = 1.597 | 0.1087 |
| Repeated EtOH: C57BL/6J male vs Gabrd^-/-^ male (Day 5 - within base) | Figure 5C | Frequency x Genotype | 5.249 | 4 | 1.312 | F (4, 48) = 7.639 | 0.0001 |
|  |  | Frequency | 3.494 | 4 | 0.8734 | F (1.432, 17.19) = 5.084 | 0.027 |
|  |  | Genotype | 0.7873 | 1 | 0.7873 | F (1, 12) = 3.040 | 0.1068 |
|  |  | Subject | 3.108 | 12 | 0.259 | F (12, 48) = 1.508 | 0.1543 |
| Repeated EtOH: C57BL/6J male vs Gabrd^-/-^ male (Day 5 - from base) | Figure 5D | Frequency x genotype | 3.504 | 4 | 0.8759 | F (4, 48) = 7.116 | 0.0001 |
|  |  | Frequency | 7.927 | 4 | 1.982 | F (1.315, 15.78) = 16.10 | 0.0005 |
|  |  | genotype | 1.396 | 1 | 1.396 | F (1, 12) = 6.071 | 0.0298 |
|  |  | Subject | 2.759 | 12 | 0.2299 | F (12, 48) = 1.868 | 0.0632 |
| Repeated EtOH: C57BL/6J female (Acute vs Day 2) | Figure 6B | Frequency x Day | 0.1412 | 4 | 0.0353 | F (4, 64) = 0.6203 | 0.6497 |
|  |  | Frequency | 1.712 | 4 | 0.428 | F (1.298, 20.76) = 7.520 | 0.0081 |
|  |  | Day | 0.03213 | 1 | 0.03213 | F (1, 16) = 0.8681 | 0.3653 |
|  |  | Subject | 0.5923 | 16 | 0.03702 | F (16, 64) = 0.6505 | 0.8298 |
| Repeated EtOH: C57BL/6J female (Day 2 vs Day 5 - within base) | Figure 6C | Frequency x Day | 0.01275 | 4 | 0.00319 | F (4, 64) = 0.04663 | 0.9958 |
|  |  | Frequency | 0.4931 | 4 | 0.1233 | F (1.285, 20.56) = 1.803 | 0.1951 |
|  |  | Day | 0.00195 | 1 | 0.00195 | F (1, 16) = 0.07268 | 0.7909 |
|  |  | Subject | 0.4284 | 16 | 0.02677 | F (16, 64) = 0.3916 | 0.98 |
| Repeated EtOH: C57BL/6J female (Day 2 vs Day 5 - from base) | Figure 6D | Frequency x Day | 0.00446 | 4 | 0.00112 | F (4, 64) = 0.01388 | 0.9996 |
|  |  | Frequency | 2.2 | 4 | 0.55 | F (1.241, 19.86) = 6.843 | 0.0123 |
|  |  | Day | 0.05616 | 1 | 0.05616 | F (1, 16) = 1.884 | 0.1888 |
|  |  | Subject | 0.477 | 16 | 0.02981 | F (16, 64) = 0.3709 | 0.9848 |
| Repeated EtOH: Gabrd^-/-^ female (Day 2 vs Day 5 - within base) | Figure 6E | Frequency x Day | 0.0591 | 4 | 0.01478 | F (4, 52) = 0.1485 | 0.9629 |
|  |  | Frequency | 0.09739 | 4 | 0.02435 | F (1.279, 16.63) = 0.2446 | 0.6856 |
|  |  | Day | 0.02673 | 1 | 0.02673 | F (1, 13) = 1.122 | 0.3087 |
| Repeated EtOH: Gabrd^-/-^ female (Day 2 vs Day 5 - from base) | Figure 6F | Frequency x Day | 0.06198 | 4 | 0.0155 | F (4, 52) = 0.3276 | 0.8582 |
|  |  | Frequency | 1.138 | 4 | 0.2845 | F (1.504, 19.55) = 6.014 | 0.0143 |
|  |  | Day | 0.00911 | 1 | 0.00911 | F (1, 13) = 0.2963 | 0.5954 |
|  |  | Subject | 0.3996 | 13 | 0.03074 | F (13, 52) = 0.6499 | 0.8004 |
| Repeated EtOH: Gabrd^-/-^ female (Acute vs Day 2) | Ext. Fig. 6-1A | Frequency | 0.5748 | 4 | 0.1437 | F (4, 28) = 1.782 | 0.1606 |
|  |  | Day | 0.5922 | 1 | 0.5922 | F (1, 7) = 6.119 | 0.0426 |
|  |  | Frequency x Day | 1.635 | 4 | 0.4088 | F (4, 28) = 4.252 | 0.0082 |
| Repeated EtOH: Gabrd^-/-^ female (Acute vs Day 5) | Ext. Fig. 6-1B | Frequency x Day | 1.628 | 4 | 0.4069 | F (4, 52) = 3.800 | 0.0087 |
|  |  | Frequency | 0.6174 | 4 | 0.1543 | F (1.211, 15.74) = 1.441 | 0.2547 |
|  |  | Day | 0.3319 | 1 | 0.3319 | F (1, 13) = 3.481 | 0.0848 |
|  |  | Subject | 1.24 | 13 | 0.09535 | F (13, 52) = 0.8905 | 0.5677 |
| Repeated VEH: C57BL/6J female (Day 2 vs Day 5 - within base) | Ext. Fig. 6-2A | Frequency x Day | 0.3045 | 4 | 0.07613 | F (4, 48) = 2.891 | 0.0318 |
|  |  | Frequency | 0.05301 | 4 | 0.01325 | F (1.210, 14.51) = 0.5033 | 0.5237 |
|  |  | Day | 3.387 | 1 | 3.387 | F (1, 12) = 5.227 | 0.0412 |
|  |  | Subject | 7.777 | 12 | 0.6481 | F (12, 48) = 24.61 | 0.0001 |
| Repeated VEH: C57BL/6J female (Day 2 vs Day 5 - from base) | Ext. Fig. 6-2B | Frequency x Day | 0.1108 | 4 | 0.0277 | F (4, 48) = 0.3122 | 0.8684 |
|  |  | Frequency | 9.913 | 4 | 2.478 | F (1.200, 14.40) = 27.93 | 0.0001 |
|  |  | Day | 0.06332 | 1 | 0.06332 | F (1, 12) = 0.3756 | 0.5514 |
|  |  | Subject | 2.023 | 12 | 0.1686 | F (12, 48) = 1.900 | 0.0583 |
| Repeated VEH: Gabrd^-/-^ female (Day 2 vs Day 5 - within base) | Ext. Fig. 6-2C | Frequency | 0.1451 | 4 | 0.03627 | F (1.316, 7.898) = 0.2415 | 0.7006 |
|  |  | Day | 0.4467 | 1 | 0.4467 | F (1.000, 6.000) = 3.235 | 0.1222 |
|  |  | Frequency x Day | 1.818 | 4 | 0.4544 | F (1.220, 7.319) = 1.884 | 0.2145 |
| Repeated VEH: Gabrd^-/-^ female (Day 2 vs Day 5 - from base) | Ext. Fig. 6-2D | Frequency | 4.97 | 4 | 1.243 | F (1.360, 8.158) = 5.434 | 0.0402 |
|  |  | Day | 0.02101 | 1 | 0.02101 | F (1.000, 6.000) = 0.03566 | 0.8564 |
|  |  | Frequency x Day | 0.3225 | 4 | 0.08062 | F (1.426, 8.554) = 0.4528 | 0.5868 |
| Repeated EtOH: C57BL/6J female vs Gabrd^-/-^ female (Day 2 - within base) | Ext. Fig. 6-3A | Frequency x Genotype | 0.2293 | 4 | 0.05732 | F (4, 64) = 0.8883 | 0.4762 |
|  |  | Frequency | 0.05668 | 4 | 0.01417 | F (1.191, 19.06) = 0.2196 | 0.6871 |
|  |  | Genotype | 0.00743 | 1 | 0.00743 | F (1, 16) = 0.4227 | 0.5248 |
|  |  | Subject | 0.281 | 16 | 0.01756 | F (16, 64) = 0.2722 | 0.9972 |
| Repeated EtOH: C57BL/6J female vs Gabrd^-/-^ female (Day 5 - within base) | Ext. Fig. 6-3C | Frequency x Genotype | 0.2965 | 4 | 0.07413 | F (4, 52) = 0.7111 | 0.588 |
|  |  | Frequency | 0.04463 | 4 | 0.01116 | F (1.376, 17.89) = 0.1070 | 0.8246 |
|  |  | Genotype | 0.08004 | 1 | 0.08004 | F (1, 13) = 2.277 | 0.1553 |
|  |  | Subject | 0.457 | 13 | 0.03516 | F (13, 52) = 0.3372 | 0.9823 |
| Repeated EtOH: C57BL/6J female vs Gabrd^-/-^ female (Day 2 - from base) | Ext. Fig. 6-3B | Frequency x Genotype | 0.1127 | 4 | 0.02817 | F (4, 64) = 0.4427 | 0.7773 |
|  |  | Frequency | 1.507 | 4 | 0.3768 | F (1.336, 21.37) = 5.922 | 0.0167 |
|  |  | Genotype | 0.00966 | 1 | 0.00966 | F (1, 16) = 0.2190 | 0.6461 |
|  |  | Subject | 0.7061 | 16 | 0.04413 | F (16, 64) = 0.6936 | 0.7897 |
| Repeated EtOH: C57BL/6J female vs Gabrd^-/-^ female (Day 5 - from base) | Ext. Fig. 6-3D | Frequency x Genotype | 0.03797 | 4 | 0.00949 | F (4, 52) = 0.1398 | 0.9667 |
|  |  | Frequency | 1.653 | 4 | 0.4132 | F (1.299, 16.89) = 6.084 | 0.0184 |
|  |  | Genotype | 0.04488 | 1 | 0.04488 | F (1, 13) = 3.421 | 0.0872 |
|  |  | Subject | 0.1706 | 13 | 0.01312 | F (13, 52) = 0.1932 | 0.9988 |
| Repeated EtoH: C57BL/6J male vs female (First day - within base) | Figure 7A | Frequency x Sex | 1.657 | 4 | 0.4142 | F (4, 64) = 4.164 | 0.0046 |
|  |  | Frequency | 0.3625 | 4 | 0.09061 | F (1.400, 22.40) = 0.9108 | 0.3834 |
|  |  | Sex | 0.4269 | 1 | 0.4269 | F (1, 16) = 5.658 | 0.0302 |
|  |  | Subject | 1.207 | 16 | 0.07546 | F (16, 64) = 0.7585 | 0.7241 |
| Repeated EtOH: C57BL/6J male vs female (First day - from base) | Figure 7B | Frequency x Sex | 0.03151 | 4 | 0.00788 | F (4, 64) = 0.06856 | 0.9912 |
|  |  | Frequency | 2.258 | 4 | 0.5645 | F (1.278, 20.44) = 4.912 | 0.0305 |
|  |  | Sex | 0.3755 | 1 | 0.3755 | F (1, 16) = 2.435 | 0.1382 |
|  |  | Subject | 2.467 | 16 | 0.1542 | F (16, 64) = 1.342 | 0.2007 |
| Repeated EtOH: C57BL/6J male vs female (Last day - within base) | Figure 7C | Frequency x Sex | 5.683 | 4 | 1.421 | F (4, 48) = 8.045 | 0.0001 |
|  |  | Frequency | 3.17 | 4 | 0.7925 | F (1.487, 17.84) = 4.488 | 0.0355 |
|  |  | Sex | 1.019 | 1 | 1.019 | F (1, 12) = 3.740 | 0.077 |
|  |  | Subject | 3.268 | 12 | 0.2723 | F (12, 48) = 1.542 | 0.1422 |
| Repeated EtOH: C57BL/6J male vs female (Last day - from base) | Figure 7D | Frequency x Sex | 3.289 | 4 | 0.8221 | F (4, 48) = 5.640 | 0.0008 |
|  |  | Frequency | 8.294 | 4 | 2.074 | F (1.351, 16.22) = 14.23 | 0.0008 |
|  |  | Sex | 2.05 | 1 | 2.05 | F (1, 12) = 9.412 | 0.0098 |
|  |  | Subject | 2.614 | 12 | 0.2178 | F (12, 48) = 1.495 | 0.1593 |

**Table 4-2.** Summary of Šídák's multiple comparisons tests for repeated alcohol experiments.

| **Description** | **Figure** | **Comparisons** | **P value** | **Mean Diff** | **SEM** | ***n*** |
| --- | --- | --- | --- | --- | --- | --- |
| Repeated EtOH: C57BL/6J male (Day 1 vs Day 5 - within base) | Figure 4C | 2-5 | 0.9599 | -0.4253 | 0.5649 | 8,6 |
|  |  | 6-12 | 0.3997 | -0.3542 | 0.1928 | 8,6 |
|  |  | 15-30 | 0.6534 | -0.2552 | 0.1777 | 8,6 |
|  |  | 40-70 | 0.0252 | 0.3381 | 0.09726 | 8,6 |
|  |  | 80-120 | 0.0775 | 0.2176 | 0.07605 | 8,6 |
| Repeated EtOH: C57BL/6J male (Day 1 vs Day 5 - from base) | Figure 4D | 2-5 | 0.5855 | 0.8153 | 0.5326 | 8,6 |
|  |  | 6-12 | 0.3494 | 0.3636 | 0.1908 | 8,6 |
|  |  | 15-30 | 0.4527 | 0.2634 | 0.1495 | 8,6 |
|  |  | 40-70 | 0.0305 | -0.3707 | 0.1051 | 8,6 |
|  |  | 80-120 | 0.0635 | -0.2437 | 0.07769 | 8,6 |
| Repeated EtOH: Gabrd^-/-^ male (Day 1 vs Day 5 - within base) | Figure 4E | 2-5 | 0.8036 | 0.2003 | 0.1702 | 8,8 |
|  |  | 6-12 | 0.6379 | 0.1878 | 0.1274 | 8,8 |
|  |  | 15-30 | 0.4217 | 0.1254 | 0.06709 | 8,8 |
|  |  | 40-70 | 0.9594 | -0.145 | 0.1913 | 8,8 |
|  |  | 80-120 | 0.9872 | -0.05647 | 0.09781 | 8,8 |
| Repeated EtOH: Gabrd^-/-^ male (Day 1 vs Day 5 - from base) | Figure 4F | 2-5 | 0.351 | -0.2936 | 0.1452 | 8,8 |
|  |  | 6-12 | 0.8943 | -0.08774 | 0.08999 | 8,8 |
|  |  | 15-30 | 0.724 | -0.06599 | 0.04983 | 8,8 |
|  |  | 40-70 | 0.0728 | 0.3224 | 0.1006 | 8,8 |
|  |  | 80-120 | 0.1669 | 0.1496 | 0.05771 | 8,8 |
| Repeated VEH: C57BL/6J male (Day 1 vs Day 5 - within base) | Ext. Fig. 4-1A | 2-5 | 0.9527 | -0.4885 | 0.5908 | 4,3 |
|  |  | 6-12 | 0.9698 | -0.1231 | 0.1633 | 4,3 |
|  |  | 15-30 | 0.9827 | -0.07075 | 0.1095 | 4,3 |
|  |  | 40-70 | 0.6576 | 0.2858 | 0.177 | 4,3 |
|  |  | 80-120 | 0.5441 | 0.1522 | 0.08182 | 4,3 |
| Repeated VEH: C57BL/6J male (Day 1 vs Day 5 - from base) | Ext. Fig. 4-1B | 2-5 | >0.9999 | 0.01558 | 0.7971 | 4,3 |
|  |  | 6-12 | 0.9176 | -0.2963 | 0.3168 | 4,3 |
|  |  | 15-30 | 0.9048 | -0.146 | 0.1493 | 4,3 |
|  |  | 40-70 | >0.9999 | 0.05855 | 0.2738 | 4,3 |
|  |  | 80-120 | 0.9837 | 0.1147 | 0.1827 | 4,3 |
| Repeated VEH: Gabrd^-/-^ male (Day 1 vs Day 5 - within base) | Ext. Fig. 4-1C | 2-5 | 0.9929 | -0.1177 | 0.219 | 4,4 |
|  |  | 6-12 | 0.9972 | 0.02187 | 0.05006 | 4,4 |
|  |  | 15-30 | 0.8109 | 0.03624 | 0.02779 | 4,4 |
|  |  | 40-70 | 0.9671 | -0.1189 | 0.1534 | 4,4 |
|  |  | 80-120 | 0.9996 | -0.01895 | 0.06603 | 4,4 |
| Repeated VEH: Gabrd^-/-^ male (Day 1 vs Day 5 - from base) | Ext. Fig. 4-1D | 2-5 | >0.9999 | -0.02151 | 0.1799 | 4,4 |
|  |  | 6-12 | 0.9445 | -0.1341 | 0.1506 | 4,4 |
|  |  | 15-30 | 0.8763 | -0.08277 | 0.07343 | 4,4 |
|  |  | 40-70 | 0.9817 | -0.391 | 0.5832 | 4,4 |
|  |  | 80-120 | 0.9966 | -0.04641 | 0.102 | 4,4 |
| Repeated EtOH: C57BL/6J male vs Gabrd^-/-^ male (Day 1 - within base) | Figure 5A | 2-5 | 0.8439 | 0.3076 | 0.2887 | 8,8 |
|  |  | 6-12 | 0.9994 | 0.04274 | 0.1438 | 8,8 |
|  |  | 15-30 | 0.9992 | 0.02869 | 0.0917 | 8,8 |
|  |  | 40-70 | >0.9999 | -0.0293 | 0.1553 | 8,8 |
|  |  | 80-120 | 0.959 | -0.06915 | 0.093 | 8,8 |
| Repeated EtOH: C57BL/6J male vs Gabrd^-/-^ male (Day 1 - from base) | Figure 5B | 2-5 | 0.6654 | -0.4309 | 0.3118 | 8,8 |
|  |  | 6-12 | 0.0138 | -0.5848 | 0.1574 | 8,8 |
|  |  | 15-30 | 0.0022 | -0.4426 | 0.08938 | 8,8 |
|  |  | 40-70 | >0.9999 | -0.02834 | 0.1635 | 8,8 |
|  |  | 80-120 | >0.9999 | 0.012 | 0.1085 | 8,8 |
| Repeated EtOH: C57BL/6J male vs Gabrd^-/-^ male (Day 5 - within base) | Figure 5C | 2-5 | 0.4847 | 0.9331 | 0.5093 | 6,8 |
|  |  | 6-12 | 0.0553 | 0.5848 | 0.1778 | 6,8 |
|  |  | 15-30 | 0.221 | 0.4093 | 0.1662 | 6,8 |
|  |  | 40-70 | 0.0343 | -0.5125 | 0.1466 | 6,8 |
|  |  | 80-120 | 0.0062 | -0.3432 | 0.0806 | 6,8 |
| Repeated EtOH: C57BL/6J male vs Gabrd^-/-^ male (Day 5 - from base) | Figure 5D | 2-5 | 0.3689 | -0.9598 | 0.4605 | 6,8 |
|  |  | 6-12 | 0.0691 | -0.4991 | 0.1495 | 6,8 |
|  |  | 15-30 | 0.118 | -0.3986 | 0.1322 | 6,8 |
|  |  | 40-70 | 0.2074 | 0.2747 | 0.1178 | 6,8 |
|  |  | 80-120 | 0.0203 | 0.156 | 0.04355 | 6,8 |
| Repeated EtOH: C57BL/6J female (Acute vs Day 2) | Figure 6B | 2-5 | 0.9378 | 0.0917 | 0.112 | 8,10 |
|  |  | 6-12 | 0.9279 | 0.07371 | 0.08668 | 8,10 |
|  |  | 15-30 | 0.269 | 0.1243 | 0.05969 | 8,10 |
|  |  | 40-70 | 0.9692 | -0.09853 | 0.1433 | 8,10 |
|  |  | 80-120 | >0.9999 | -0.001044 | 0.1204 | 8,10 |
| Repeated EtOH: C57BL/6J female (Day 2 vs Day 5 - within base) | Figure 6C | 2-5 | >0.9999 | 0.01583 | 0.08785 | 10,8 |
|  |  | 6-12 | >0.9999 | 0.002994 | 0.08448 | 10,8 |
|  |  | 15-30 | >0.9999 | -0.001355 | 0.04361 | 10,8 |
|  |  | 40-70 | >0.9999 | -0.0215 | 0.213 | 10,8 |
|  |  | 80-120 | 0.9859 | 0.05083 | 0.08812 | 10,8 |
| Repeated EtOH: C57BL/6J female (Day 2 vs Day 5 - from base) | Figure 6D | 2-5 | 0.9946 | -0.05899 | 0.1261 | 10,8 |
|  |  | 6-12 | 0.9824 | -0.05479 | 0.09024 | 10,8 |
|  |  | 15-30 | 0.9106 | -0.05266 | 0.05819 | 10,8 |
|  |  | 40-70 | 0.9989 | -0.06221 | 0.1888 | 10,8 |
|  |  | 80-120 | >0.9999 | -0.02272 | 0.1225 | 10,8 |
| Repeated EtOH: Gabrd^-/-^ female (Day 2 vs Day 5 - within base) | Figure 6E | 2-5 | 0.9961 | -0.1036 | 0.2356 | 8,7 |
|  |  | 6-12 | >0.9999 | -0.01972 | 0.117 | 8,7 |
|  |  | 15-30 | 0.9905 | 0.04232 | 0.08003 | 8,7 |
|  |  | 40-70 | >0.9999 | -0.008155 | 0.1943 | 8,7 |
|  |  | 80-120 | 0.641 | -0.1 | 0.07087 | 8,7 |
| Repeated EtOH: Gabrd^-/-^ female (Day 2 vs Day 5 - from base) | Figure 6F | 2-5 | 0.9999 | 0.03155 | 0.1489 | 8,7 |
|  |  | 6-12 | 0.9937 | 0.04228 | 0.08741 | 8,7 |
|  |  | 15-30 | 0.9977 | -0.02897 | 0.07408 | 8,7 |
|  |  | 40-70 | 0.9395 | -0.1186 | 0.1454 | 8,7 |
|  |  | 80-120 | 0.9708 | -0.03667 | 0.05355 | 8,7 |
| Repeated EtOH: Gabrd^-/-^ female (Acute vs Day 2) | Ext. Fig. 6-1A | 2-5 | 0.0019 | 0.6267 | 0.155 | 8,8 |
|  |  | 6-12 | 0.4993 | 0.2424 | 0.155 | 8,8 |
|  |  | 15-30 | 0.4732 | 0.2484 | 0.155 | 8,8 |
|  |  | 40-70 | 0.6825 | -0.2011 | 0.155 | 8,8 |
|  |  | 80-120 | 0.9983 | -0.05599 | 0.155 | 8,8 |
| Repeated EtOH: Gabrd^-/-^ female (Acute vs Day 5) | Ext. Fig. 6-1B | 2-5 | 0.3337 | 0.5728 | 0.2899 | 8,7 |
|  |  | 6-12 | 0.1189 | 0.2672 | 0.1046 | 8,7 |
|  |  | 15-30 | 0.15 | 0.2083 | 0.08645 | 8,7 |
|  |  | 40-70 | 0.4916 | -0.2586 | 0.1559 | 8,7 |
|  |  | 80-120 | 0.3951 | -0.1231 | 0.06824 | 8,7 |
| Repeated VEH: C57BL/6J female (Day 2 vs Day 5 - within base) | Ext. Fig. 6-2A | 2-5 | 0.8631 | -0.2987 | 0.2802 | 8,6 |
|  |  | 6-12 | 0.7917 | -0.3186 | 0.262 | 8,6 |
|  |  | 15-30 | 0.5515 | -0.4104 | 0.2429 | 8,6 |
|  |  | 40-70 | 0.0612 | -0.6408 | 0.1878 | 8,6 |
|  |  | 80-120 | 0.1886 | -0.554 | 0.2057 | 8,6 |
| Repeated VEH: C57BL/6J female (Day 2 vs Day 5 - from base) | Ext. Fig. 6-2B | 2-5 | 0.6031 | -0.1919 | 0.129 | 8,6 |
|  |  | 6-12 | 0.8273 | -0.09669 | 0.08775 | 8,6 |
|  |  | 15-30 | 0.9383 | -0.05288 | 0.06362 | 8,6 |
|  |  | 40-70 | >0.9999 | 0.04238 | 0.3096 | 8,6 |
|  |  | 80-120 | >0.9999 | -0.004796 | 0.1152 | 8,6 |
| Repeated VEH: Gabrd^-/-^ female (Day 2 vs Day 5 - with in base) | Ext. Fig. 6-2C | 2-5 | 0.5662 | 0.6214 | 0.3808 | 7,7 |
|  |  | 6-12 | 0.616 | 0.3371 | 0.2187 | 7,7 |
|  |  | 15-30 | 0.3124 | 0.2325 | 0.1067 | 7,7 |
|  |  | 40-70 | 0.9116 | -0.2865 | 0.3054 | 7,7 |
|  |  | 80-120 | 0.9527 | -0.1057 | 0.1329 | 7,7 |
| Repeated VEH: Gabrd^-/-^ female (Day 2 vs Day 5 - from base) | Ext. Fig. 6-2D | 2-5 | 0.9994 | -0.1318 | 0.4359 | 7,7 |
|  |  | 6-12 | 0.9981 | 0.1007 | 0.2622 | 7,7 |
|  |  | 15-30 | 0.962 | 0.1293 | 0.1718 | 7,7 |
|  |  | 40-70 | 0.9003 | -0.2281 | 0.2351 | 7,7 |
|  |  | 80-120 | 0.9997 | -0.04342 | 0.1682 | 7,7 |
| Repeated EtOH: C57BL/6J female vs Gabrd^-/-^ female (Day 2 - within base) | Ext. Fig. 6-3A | 2-5 | 0.8165 | -0.144 | 0.1295 | 10,8 |
|  |  | 6-12 | 0.9257 | -0.08176 | 0.09511 | 10,8 |
|  |  | 15-30 | 0.8923 | -0.06936 | 0.07249 | 10,8 |
|  |  | 40-70 | 0.9593 | 0.1138 | 0.1547 | 10,8 |
|  |  | 80-120 | 0.8643 | 0.08986 | 0.08875 | 10,8 |
| Repeated EtOH: C57BL/6J female vs Gabrd^-/-^ female (Day 2 - from base) | Ext. Fig. 6-3B | 2-5 | 0.9988 | 0.03872 | 0.114 | 10,8 |
|  |  | 6-12 | 0.7103 | -0.09834 | 0.07682 | 10,8 |
|  |  | 15-30 | >0.9999 | -0.008148 | 0.05969 | 10,8 |
|  |  | 40-70 | 0.969 | 0.1133 | 0.1645 | 10,8 |
|  |  | 80-120 | 0.9924 | 0.05872 | 0.1163 | 10,8 |
| Repeated EtOH: C57BL/6J female vs Gabrd^-/-^ female (Day 5 - within base) | Ext. Fig. 6-3C | 2-5 | 0.7814 | -0.2634 | 0.2155 | 8,7 |
|  |  | 6-12 | 0.8889 | -0.1045 | 0.1085 | 8,7 |
|  |  | 15-30 | 0.9949 | -0.02568 | 0.05525 | 8,7 |
|  |  | 40-70 | 0.991 | 0.1272 | 0.2433 | 8,7 |
|  |  | 80-120 | 0.924 | -0.061 | 0.07008 | 8,7 |
| Repeated EtOH: C57BL/6J female vs Gabrd^-/-^ female (Day 5 - from base) | Ext. Fig. 6-3D | 2-5 | 0.9397 | 0.1293 | 0.1583 | 8,7 |
|  |  | 6-12 | >0.9999 | -0.001273 | 0.09941 | 8,7 |
|  |  | 15-30 | 0.9999 | 0.01553 | 0.07287 | 8,7 |
|  |  | 40-70 | 0.999 | 0.05689 | 0.1724 | 8,7 |
|  |  | 80-120 | 0.9725 | 0.04477 | 0.06596 | 8,7 |
| Repeated EtoH: C57BL/6J male vs female (First day - within base) | Figure 7A | 2-5 | 0.2719 | 0.573 | 0.2639 | 8,10 |
|  |  | 6-12 | 0.2805 | 0.2601 | 0.1252 | 8,10 |
|  |  | 15-30 | 0.3978 | 0.1615 | 0.08753 | 8,10 |
|  |  | 40-70 | 0.6836 | -0.1765 | 0.1337 | 8,10 |
|  |  | 80-120 | 0.6639 | -0.1251 | 0.09227 | 8,10 |
| Repeated EtOH: C57BL/6J male vs female (First day - from base) | Figure 7B | 2-5 | 0.9929 | -0.146 | 0.2898 | 8,10 |
|  |  | 6-12 | 0.8999 | -0.1319 | 0.1396 | 8,10 |
|  |  | 15-30 | 0.7422 | -0.1062 | 0.08405 | 8,10 |
|  |  | 40-70 | 0.7706 | -0.1888 | 0.1598 | 8,10 |
|  |  | 80-120 | 0.9841 | -0.07705 | 0.1303 | 8,10 |
| Repeated EtOH: C57BL/6J male vs female (Last day - within base) | Figure 7C | 2-5 | 0.4119 | 1.014 | 0.5072 | 6,8 |
|  |  | 6-12 | 0.0435 | 0.6173 | 0.1692 | 6,8 |
|  |  | 15-30 | 0.2117 | 0.4153 | 0.1606 | 6,8 |
|  |  | 40-70 | 0.112 | -0.5361 | 0.1922 | 6,8 |
|  |  | 80-120 | 0.0122 | -0.2919 | 0.07094 | 6,8 |
| Repeated EtOH: C57BL/6J male vs female (Last day - from base) | Figure 7D | 2-5 | 0.3202 | -1.02 | 0.4643 | 6,8 |
|  |  | 6-12 | 0.0433 | -0.5503 | 0.1582 | 6,8 |
|  |  | 15-30 | 0.093 | -0.4222 | 0.1367 | 6,8 |
|  |  | 40-70 | 0.9418 | 0.1197 | 0.1454 | 6,8 |
|  |  | 80-120 | 0.2263 | 0.144 | 0.06377 | 6,8 |
